# Synthetic and plant-derived multivalent galactans as modulators of cancer-associated galectins-3 and -9

**DOI:** 10.1101/2025.01.14.632949

**Authors:** Lukas Pfeifer, Kim-Kristine Mueller, Maximilian Thal Müller, Lisa-Marie Philipp, Susanne Sebens, Birgit Classen

## Abstract

Galectins are β-galactoside-binding proteins with numerous functions. Some of them are involved in proliferation and metastasis of cancer, making them promising therapeutic targets. As different plant glycans have been shown to bind to galectins, plant saccharides might be potential galectin inhibitors. To produce plant galactans rich in galactose and smaller in size, we degraded arabinogalactan-proteins from *Echinacea purpurea* and *Zostera marina* as well as arabinogalactan from larch. As galectin (Gal)-3 and -9 both have been described to be involved in cancer development, we quantified the binding capacities of the different galactans to both galectins by biolayer-interferometry. Our results revealed that all plant-derived galactans and Yariv reagents with terminal galactose and lactose residues bind to Gal-3 in micromolar ranges. Surprisingly, only the higher charged galactans from *Zostera marina* showed affinity to Gal-9. Investigations of two different pancreatic cancer cell lines (Panc1 and Panc89) and different cell variants thereof revealed that Gal-3 was expressed by both cell lines with a significantly higher Gal-3 level in Panc1 cells compared to Panc89 cells. Conversely, Gal-9 was only detected in Panc89 cells. The findings revealed that galactans are promising sources to develop galectin antagonists and plant galactans from different species express specificities for distinct galectins.

## 1. Introduction

Galectins are an ancient family of β-galactoside carbohydrate binding proteins, which share at least one carbohydrate recognition domain (CRD) consisting of ∼130 amino acids of which only six amino acids are directly involved in carbohydrate binding [1,2]. The 15 mammalian galectins described can be divided into three groups: prototypical, chimeric, and tandem repeat [3]. Galectin (Gal)-1, -2, -5, -7, -10, -11, -13, -14, -15 belong to the prototypical galectins, which have a single CRD and commonly form non-covalently linked homodimers. Gal-3 is the only chimeric galectin with a single C-terminal CRD linked to a large N-terminus of about 120 – 160 amino acids, through which Gal-3 can form pentamers. Gal-4, -8, -9, -12 are tandem-repeat galectins with two different CRDs connected by a linker peptide of five to more than 50 amino acids [2] which can form multivalent lattices [4].

Galectins have a variety of roles in normal physiology including development, pattern recognition, immunity but also in pathogenesis of autoimmune diseases and cancer [3]. They are involved in proliferation and metastasis of tumor cells and are central mediators of immune escape, e.g. by promotion of T cell apoptosis and inhibition of T cell activation [5,6]. Especially Gal-1, -3 and -9 are notable for their roles in tumor immune escape and have been widely studied in the context of tumor immunology [4]. Due to this crucial involvement in tumor progression, galectins are potential targets for cancer therapy. One of the most lethal cancers with a high therapy resistance and still poor prognosis is pancreatic ductal adenocarcinoma (PDAC; [6]). Different studies have shown that Gal-3 and -9 promote immune evasion of PDAC cells, e.g. by influencing T cell differentiation and viability (for review, see [4,6]). Furthermore, binding of Gal-9 to Dectin 1 on macrophages has been shown to result in polarization into a tolerogenic macrophage phenotype suppressing the adaptive anti-tumor immunity and thereby promoting PDAC development [7]. Relative to other human solid tumors, PDAC showed the highest expression of Gal-9 mRNA expression [8]. Their expression levels were much higher than PD-L1 levels which explains the poor response of PDAC to anti-PD-1 checkpoint inhibitors. In PDAC patients, elevated levels of Gal-3 as well as Gal-9 have been detected in tumor tissues compared to pancreatic tissues from patients with benign pancreatic diseases [8–11]. Altogether, these findings support the potential of Gal-3 and -9 as tumor associated target structures which can be addressed to improve PDAC treatment. Anti-cancer therapeutic approaches targeting galectins are currently in the focus of scientific interest [4] and include poly-N-acetyllactosamine (derivatives), small molecule glycomimetics, multivalent materials decorated with glycans which mimic the composition of the glycocalyx and the extracellular matrix and finally plant-derived saccharides [12]. Some are currently in pre-clinical and clinical trials [4,13]. Today, clinical strategies to block Gal-9 are mainly based on monoclonal antibodies, but potential inhibitors of Gal-3 often include plant derived polysaccharides [4], especially modified citrus pectin [14–16] and galactomannan [17]. In contrast to many other inhibitors, pectins and galactomannans (having rather low contents of galactose) bind only weakly to the canonical galactoside-binding sites of galectins [18] but in other regions of Gal-3 [15,19]. Galactose is the dominant monosaccharide in arabinogalactans. These plant macromolecules are either present as pure polysaccharides, especially in the wood of larch and some other gymnosperms [20], or alternatively as so-called arabinogalactan-proteins (AGPs). AGPs consist of several arabinogalactan (AG) moieties (around 90 % of the molecule), which are covalently linked to a small protein backbone *via* hydroxyproline [21]. In both cases, the arabinogalactans belong to type II with a β-D-1,3-galactan core including β-D-1,3,6-galactose branching points for attachment of β-D-1,6-galactan side chains, terminating in mainly arabinose (Ara) residues [22]. We performed partial degradation of AG from larch wood as well as from AGPs of the medicinal plant *Echinacea purpurea* and the seagrass *Zostera marina* with the aim to reduce the molecular weight and relatively increase the galactose content. Whereas the AGP of *Echinacea purpurea* is a typical angiosperm land plant AGP [23], *Zostera* AGP is special with high amounts of β-D-glucuronic acid (GlcA) residues [24]. All degraded galactans are rich in β-D-1,3-galactose – surprisingly similar to the poly-β-galactosyl epitope (Galβ1-3)^n^ found on the parasite *Leishmania* which is recognized by Gal-3 and -9 [25]. Furthermore, plant β-(1,3)-galactosyltransferases have been identified which possess a galectin domain [26,27]. Therefore, due to the conserved nature of galectins, binding of these plant galactans also to human galectins seems probable. Additionally, we synthesized tri-antennary Yariv’s reagents [28] with terminal galactose and lactose residues as potential galectin inhibitors.

The aim of this study was to generate a palette of (macro-)molecules with the potential to inhibit PDAC-related Gal-3 and -9 and elucidate their binding capacities by biolayer interferometry (BLI). These two galectins are promising targets for inhibition of PDAC progression as they are found to be highly upregulated in several PDAC cell lines or biopsy samples [8,29]. To support the relevance of these galectins as potential therapeutic targets, our investigations were complemented by protein level determination of Gal-3 and -9 in six different PDAC cell variants. Inhibitory strategies of Gal-9 including plant-derived galactans are currently not described. Therefore, this study provides basic information about the potential applicability of non-toxic carbohydrate derivatives in the therapy of Gal-9 associated pathogenic processes. The results will broaden the understanding of intraction between plant derived and synthetic galactans to human Gal-3 and -9 and increase our knowledge on favorable structures of possible galectin inhibitors. Although *in-vitro* studies are not sufficient to evaluate the therapeutic effectiveness, they are mandatory to provide important basic information for future studies focusing on inhibitor design. Especially in the field of cancer types exhibiting a high therapy resistance and a poor prognosis – like PDAC – there is a deep need for new and innovative therapeutic concepts.

## 2. Material and methods

### 2.1. Plant material

Fresh rhizomes of the seagrass *Zostera marina* L. were collected from the Baltic Sea beach of Schilksee, Kiel, Germany (54°25′38.2″N 10°10′17.5″E) in September 2022. All samples were cleaned with tap water and freeze-dried (Christ Alpha 1-4 LSC, Martin Christ Gefriertrocknungsanlagen GmbH, Osterode, Germany). Pressed juice of *Echinacea purpurea* L. MOENCH was a kind gift of Madaus GmbH, Cologne, Germany. The high molecular weight fraction, here called aqueous extract (AE), was isolated according to [23]. Larch arabinogalactan (LAG) was purchased from Sigma-Aldrich Chemie GmbH, Taufkirchen, Germany.

### 2.2. Yariv reagents

Yariv reagents (1,3,5-tri(*p*-glycosyloxyphenylazo)-2,4,6-trihydroxybenzenes) with terminal α- and β-galactose, lactose and β-Glc were synthesized according to [28]. First step is the hydrogenation of *p*-nitrophenyl-(α/β)-D-glycopyranoside (Sigma-Aldrich Chemie GmbH) to *p*-aminophenyl-β-D-glycopyranoside (α-galactose, β-galactose, β-glucose). The *p*-aminophenyl-β-D-galactopyranoside (Sigma-Aldrich Chemie GmbH) and the *p*-aminophenyl-β-D-lactopyranoside (Toronto Research Chemicals, North York, Kanada) were purchased. These glycopyranosides were then converted to the diazonium salt by addition of sodium nitrite. Finally, this diazonium salt is coupled to phloroglucinole at constant alkaline pH 9 by an autotitrator (Metrohm 719 S-Titrino, Deutsche METROHM GmbH & Co. KG, Filderstadt, Germany). The red coloured product is precipitated by methanol overnight at 4 °C, separated by centrifugation (4,000 g, 20 min, Heraeus Megafuge 16, Thermo Scientific, Waltham, USA) and freeze-dried (Christ Alpha 1-4 LSC).

### 2.3. Aqueous extract (AE) from *Zostera marina* rhizomes

After four pre-extractions of the freeze-dried and ground (Silvercrest, Kompernass H. GmbH, Bochum, Germany) rhizomes with 70 % acetone solution (V/V), the plant residue was extracted with double-distilled water (ddH_2_O). Each extraction process was performed under constant stirring (SM 2484, Edmund Bühler GmbH, Bodelshausen, Germany) at 4 °C in a 1:10 (w/V) ratio for 22 h. The aqueous extract was separated by a tincture press (HAFICO HP 2 H, Fischer Tinkturenpressen GmbH, Mönchengladbach, Germany). After incubating the aqueous extract in a water bath (GFL 1002, LAUDA-GFL Gesellschaft für Labortechnik mbH, Burgwedel, Germany) at 90 – 95 °C for ten minutes, the denaturated proteins were removed by centrifugation (4,122 *g*, 20 min, 4 °C, Heraeus Multifuge X3R, Thermo Scientific, Waltham, USA). Subsequently the aqueous extract was evaporated in a rotary evaporator (40 °C, 0,010 mbar; Laborota 4000, Heidolph Instruments GmbH & CO. KG, Schwabach, Germany) to approximately one-tenth of its volume and then added to cooled ethanol in a 1:4 (V/V) ratio. Following incubation overnight at 4°C, the precipitate was isolated by centrifugation (4 °C, 4,122 g, 20 min, Heraeus Multifuge X3R) and finally freeze-dried (AE, Christ Alpha 1-4 LSC).

### 2.4. Isolation of arabinogalactan-proteins (AGPs)

The AE of *Z. marina* and *E. purpurea* were used to isolate AGPs by β-glucose-Yariv reagent (βGlcY), according to the procedure of [30]. The AE solution, dissolved in ddH_2_O, and the βGlcY solution (1 mg ml^-1^), dissolved in an equal volume of sodium chloride solution (0.3 mol l^-1^), were combined and incubated overnight at 4 °C to precipitate the AGP-βGlcY-complex. This complex was separated by centrifugation (19,000 *g*, 4 °C, 20 min, Heraeus Multifuge X3R). After re-dissolving in ddH_2_O and heating to 50°C, sodium hydrosulfite was added, until the red complex became colourless. At the end, the aqueous AGP solution was dialyzed against demineralized water (4 °C; 4 d; MWCO 14 kDa, Membra-Cel® MD25 14 x 500 CLR, Carl Roth GmbH, Karlsruhe, Germany) and freeze-dried (Christ Alpha 1-4 LSC).

### 2.5. Analysis of monosaccharides

Determination of the neutral monosaccharide composition was performed according to [31] with slight modifications [32]. To identify and quantify the neutral monosaccharides a gas chromatography (GC) with flame ionization detection (FID) and mass spectrometry detection (MSD) was used: GC + FID: 7890B; Agilent Technologies, USA; MS: 5977B MSD; Agilent Technologies, USA; column: Optima-225; Macherey-Nagel, Germany; 25 m, 250 μm, 0.25 μm; helium flow rate: 1 ml min^-1^; split ratio 30:1. A temperature gradient was implemented to achieve peak separation (initial temperature 200 °C, subsequent holding time of 3 min; final temperature 243 °C with a gradient of 2 °C min^-1^).

### 2.6. Partial degradations of AGPs and LAG

AGPs were subjected to alkaline degradation to decompose their protein backbones. AGPs were dissolved in 0.44 mol l^-1^ sodium hydroxide and incubated for 18 h at 105 °C (Bioblock Scientific 92607, Barnstead Thermolyne Corp., Dubuque, USA). After neutralization (761 Calimatic, Knick Elektronische Messgeräte GmbH & Co. KG, Berlin, Germany), the solution was concentrated (Laborota 4000) and added into cooled ethanol, resulting in a final ethanol concentration of 80 % (V/V). After precipitation overnight at 4°C, the precipitate was isolated by centrifugation (19,000*g*, 4 °C, 20 min, Heraeus Multifuge X3R). After two additional washing steps, the AG-precipitate was dissolved in ddH_2_O, dialyzed against demineralized water and freeze-dried (AGP_Alk_, Christ Alpha 1-4 LSC).

The AGP_Alk_ and the LAG samples were then subjected to oxalic acid hydrolysis to remove labile terminal furanosidic arabinose residues. Therefore, a modified method of [33] was used. The sample was dissolved in 12.5 mmol l^-1^ oxalic acid and incubated for 5 h at 100 °C (Bioblock Scientific 92607). The hydrolysate was added to cooled ethanol (final concentration of 80 % [V/V]) and incubated overnight at 4 °C. The precipitate (19,000 *g*, 4 °C, 20 min, Heraeus Multifuge X3R) was washed two times, dissolved in ddH_2_O and freeze-dried (AGP_AlkOx_, LAG_Ox_, Christ Alpha 1-4 LSC).

For LAG, another method to remove arabinoses was used additionally. First, the LAG-sample was dissolved in 0.5 mol l^-1^ trifluoroacetic acid and incubated for 2 h at 80 °C (Bioblock Scientific 92607). The isolation of the partially degraded galactans corresponds to those of the oxalic acid hydrolysis (LAG_TFA_).

### 2.7. Sulfation of LAG_TFA_

For the chemical sulfation of arabinogalactan (AG), the methodology of [34] was used. 100 mg of dried LAG_TFA_ was mixed with 1.5 g of 1-butyl-3-methylimidazolium chloride ([C_4_mim]Cl). To ensure complete melting of the [C_4_mim]Cl and dissolution of the LAG_TFA_, the mixture was stirred for 3 h at 80 °C (RETbasic, IKA-Werke GmbH & CO. KG, Staufen, Germany) under argon atmosphere.

After dropwise addition of 300 µl chlorosulfonic acid to 1 ml anhydrous pyridine under constant stirring in an ice bath (see [35]), the activated sulfating reagent was added to the polysaccharide solution. 10 mg 4-dimethylamino-pyridine (DMAP) was dissolved in 100 µl anhydrous pyridine and admitted to the solution. This was incubated for 1 h in a 30 °C water bath (GFL 1002, GFL Gesellschaft für Labortechnik mbH, Burgwedel, Germany) and subsequently neutralized with 2.5 mol l^-1^ sodium hydroxide solution (761 Calimatic). The resulting sulfated AG solution was dialyzed two times at 4 °C with 14 kDa cut off dialysis tubes (Membra-Cel® MD25 14 x 500 CLR) and freeze-dried (Christ Alpha 1-4 LSC). This freeze-dried sample is named LAG_TFA_sulf_ in the following.

### 2.8. Elemental analysis

The elemental analysis (C, H, N and S) was performed in the Chemistry Department of the University of Hamburg, Hamburg, Germany (EA3000, Eurovector, Pavia, Italy; Unicube, Elementar Analysensysteme GmbH, Langenselbold, Germany).

### 2.9. FT-IR Spectrometry

For detection of sulfate groups, infrared spectrometry was used (IRAffinity-1S system with the MIRacle 10 single reflexion ATR accessory; Shimadzu Corporation, Kyoto, Japan). Spectra were obtained in the range from 600 to 4000 cm^−1^ with the resolution of 2 cm^−1^ by use of the LabSolutions IR software Version 2.10 (Shimadzu Corporation). The spectra were preprocessed in the Orange data mining toolbox in python [36] by using Savitzky-Golay filtering and baseline correction in rubber-band mode.

### 2.10. Photometric quantification of sulfate

Quantification of sulfate content after chemical sulfation of LAG_TFA_ was performed following the protocol of Farndale *et al*. [37]. In brief, LAG_TFA_ and LAG_TFA_Sulf_ samples were prepared in 25 µg ml^-1^ ddH_2_O concentration. Fucoidan (from *F. vesiculosus*, commercially purchased from Sigma-Aldrich Chemie GmbH) and LAG (both in 0, 5, 10, 20, 30, 40, 50 µg ml^-1^ ddH_2_O concentration) were used as positive and negative controls, respectively. 50 µl of samples or standards were combined with 1.250 ml of color reagent (dimethylmethylene blue in 1 l water containing 40 mmol glycine, 40 mmol NaCl and 1 mmol HCl, pH 3.0) and absorbance was measured at 525 nm wavelength using a spectrophotometer (UVmini-1240, Shimadzu AG, Kyoto, Japan).

### 2.11. Galectins

Gal-3 and Gal-9 were purchased from BIOZOL Diagnostica Vertrieb GmbH, Eching, Germany (for details, see Table S1).

### 2.12. Biotinylation of galectins and biolayer interferometry

Preparation of recombinant human Gal-3 and -9 for biolayer interferometry measurement (BLI) was performed according to [38]. In brief, the lyophilized galectins were solubilized and the original buffers were exchanged to phosphate-buffered saline (PBS, see Table S2 for details). Galectins were then subjected to the biotinylation procedure with *N*-hydroxysuccinimid coupled biotin (NHS-Biotin-PEG4, EZ-Link, Thermo Fisher Scientific Corp.) according to the manufacturer’s recommendations [39]. The reagent was used in 1 mmol l^-1^ concentration and a molar coupling ratio of MCR = 1 was chosen. Incubation was performed for 2 h on wet ice and unbound reagent was removed by spin column desalting. For buffer exchange (see above) and desalting the Zeba Spin desalting columns (0.5 ml, Thermo Fisher Scientific Corp.) were used. Molecular binding experiments of plant galactans to Gal-3 and -9 were performed by BLI (Octet RED96e; FortéBio, Sartorius AG, Göttingen, Germany) with Dip & Read SSA sensors (FortéBio, Sartorius AG). For each galactan sample (degraded *Echinacea*- and *Zostera* AGP, degraded larch AG as well as sulfated degraded larch AG, Yariv reagents) an array of concentrations was chosen and serial dilution was conducted. A detailed explanation of the binding experiment is given in [38] and all relevant steps are stated in Table S3. In each BLI experiment, an empty sensor was dipped into buffer solution instead of galectin and blocked subsequently with biocytin. The highest concentration of sample was tested with this control sensor to rule out unspecific binding. Afterwards, these control sensors without loaded galectin were substracted from the other concentrations. All sensorgrams were baseline corrected and an inter-step correction to the association was performed. Raw BLI sensorgrams are given in Supplementary Data S1. K_D_-values were calculated using customized steady-state analysis in SigmaPlot (version 14.5, Systat Software GmbH, Düsseldorf, Germany).

### 2.13. Cell lines and cell culture

Different PDAC cell lines (Panc1 and Panc89, see Table S4) were tested to model the characteristics of mesenchymal-like (Panc1) and epithelial (Panc89) PDAC cells. Additionally, Holo- and Paraclone cell variants (isolated and generated via single cell cloning [40,41]) of both parental Panc1 and Panc89 cells were comparatively analyzed. Holoclones are referred to as cancer stem cell fraction, Paraclones to be the more differentiated cancer cell fraction of parental PDAC cell lines [42]. All cell lines were cultivated in Panc-medium (see Table S2).

### 2.14. Western Blot analysis

For the protein level verification, a western blot analysis was performed. After cultivation, all cell variants were prepared for protein isolation. Therefore, cells were detached and centrifuged at 300 g for 5 min. Afterwards, the cell pellets were resuspended in cell lysis buffer (see Table S2). The resulting cell suspensions were sonicated with ultrasound (5×1 sec at maximum intensity; Sonicator, MSE, Nuaille, FR) on ice and centrifuged (300 g, 5 min, 4 °C) to remove cell debris. The protein containing supernatants were transferred into fresh 1.5 ml centrifugation tubes. Protein concentration in supernatants was analyzed with the BioRad DC protein Assay Kit II (500-0112, Bio-Rad Laboratories GmbH, Feldkirchen, Germany). Afterwards, 20 µg protein of each cell sample was mixed with loading dye (see Table S2), heated at 95 °C for 5 min, cooled on ice, centrifuged (300 g, short spin) and separated by SDS-PAGE (12 % polyacrylamide gel). SDS-PAGE was performed at 125 V for 1.5 h (Mini-PROTEAN Tetra Vertical Electrophoresis Cell, BioRad, Hercules, US) followed by the western blot analysis in a wet-blot chamber system (Mini Trans-Blot^®^ cell, BioRad, Hercules, US) to transfer the proteins to a transfer membrane (Immobilon P PVDF membrane, Sigma-Aldrich Chemie GmbH, Taufkirchen, Germany). Western blotting was performed at 100 V for 1.5 h, followed by membrane blocking with 25 ml 5 % milk in TBST (TRIS-buffered saline + Tween 20, see Table S2) for 1 h at room temperature (RT). After blocking, the membrane was washed 3×5 min with TBST on a shaker. Incubation of the primary antibodies against galectins and the reference protein β-actin, respectively (see details in Table S5), was performed with a dilution of 1:500 or 1:10.000, respectively, in 5 % milk in TBST at 4°C overnight on a roller. After a washing step with TBST (3×5 min on a shaker) the membrane was incubated with the related horseradish peroxidase-conjugated secondary antibodies (1:2000 in 5 % milk in TBST; see details in Table S5) for 1 h at RT on a roller. After 1 h, the membrane was washed 3×5 min with TBST on a shaker. Detection of proteins was performed by 2 min incubation with the Radiance Chemiluminescence Substrate (Azure Biosystems Inc., Dublin, CA, USA) and the western blot imaging system Azure 300Q (Azure Biosystems Inc.). The raw images (see Supplementary Data S2) were then used for quantification of relative intensity by using ImageJ 1.54g [43] with the build-in gel quantification tool.

### 2.15. Statistical analysis

Data from protein quantification were tested for normal distribution by using the Shapiro-Wilk normality test [44] using R (version 4.3.0, results are given in Table S6). One-way ANOVA test (as implemented in the R package “rstatix” [45]) was used to evaluate global *p*-value. Subsequently, all groups were subjected to pairwise comparison using t-test [46] with Holm-Bonferroni correction [47] to counteract familywise error rates (as implemented in the R package “rstatix” [45]. Results are given in Tables S7-S9.

### 2.16 Material availability

Synthesized galactan material as well as Yariv-reagents are available in limited amounts after request by the corresponding author, Birgit Classen.

## 3. Results

### 3.1. Analytical characterization of galactans

AGPs from the medicinal plant *Echinacea purpurea* and from the Baltic seagrass *Zostera marina* were partially degraded by alkaline and subsequent acid treatment. The protein moiety is destroyed by alkaline treatment (“Alk”), and mild acid hydrolysis with oxalic acid (“Ox”) cleaves labile monosaccharide linkage types, in case of AGPs mainly furanosidic Ara residues, which are located at the periphery of the molecules. The monosaccharide composition of the native samples and their partial degradation products are shown in Tables 1-3 and representative GC-chromatograms are shown in Figure S1. As expected, the monosaccharide composition of the alkaline degraded AGP samples was nearly unchanged in comparison to the native AGPs as only the protein moieties were removed (Tables 1 and 2). After partial acid hydrolysis by mild oxalic acid, all samples show an increase in galactose due to loss of Ara (Tables 1-3). For larch AG, we also performed a mild acid hydrolysis by diluted trifluoroacetic acid (TFA), which was even more effective in cleaving furanosidic Ara (Table 3). Sulfation of LAG (Sulf) did not alter the monosaccharide composition (Table 3). Highest galactose content was achieved in the larch sample LAG_TFA_ (95.3 %), followed by AGP_Alk_Ox_ from *Echinacea* (83 %) and finally AGP_Alk_Ox_ from *Zostera* (64.5 %).

**Table 1.** Neutral monosaccharide composition of the AGP and degraded AGPs (AGP_Alk_ and AGP_Alk_Ox_) of *Echinacea purpurea* in % (mol mol^-1^; n=3) determined by derivatization with following GC-FID/MS quantification as alditol acetates.

| Neutral monosaccharide | AGP | AGP<sub>Alk</sub> | AGP<sub>AlkOx</sub> |
| --- | --- | --- | --- |
| rhamnose | 1.0 ± 0.1 | 1.1 ± 0.0 | tr |
| fucose | tr | tr | - |
| arabinose | 30.1 ± 1.1 | 31.9 ± 0.6 | 15.1 ± 2.5 |
| xylose | - | - | - |
| mannose | tr | tr | tr |
| galactose | 65.4 ± 1.4 | 64.5 ± 0.3 | 83.0 ± 2.2 |
| glucose | $3.5 \pm 1.0$ | $2.5 \pm 0.4$ | $1.9 \pm 1.7$ |
| tr: trace value < 1% |  |  |  |

**Table 2.** Neutral monosaccharide composition of the AE, AGP and degraded AGPs (AGP_Alk_ and AGP_Alk_Ox_) of *Zostera marina* rhizomes in % (mol mol^-1^; n=3). The composition was determined after aldose reduction and *per*-acetylation by GC-FID/MS measurements.

| Neutral monosaccharide | AGP | AGP<sub>Alk</sub> | AGP<sub>Alk_Ox</sub> |
| --- | --- | --- | --- |
| rhamnose | $8.3 \pm 0.2$ | $7.5 \pm 0.1$ | $7.0 \pm 0.3$ |
| fucose | tr | - | - |
| arabinose | $34.1 \pm 0.7$ | $34.7 \pm 0.7$ | $23.0 \pm 0.7$ |
| xylose | tr | tr | - |
| mannose | $2.1 \pm 0.1$ | $1.5 \pm 0.0$ | tr |
| galactose | $47.4 \pm 0.9$ | $51.7 \pm 1.1$ | $64.5 \pm 1.8$ |
| glucose | $8.1 \pm 0.6$ | $4.6 \pm 0.6$ | $5.5 \pm 1.3$ |

tr: trace value < 1%

**Table 3.** Neutral monosaccharide composition of the LAG and degraded LAG (LAG_Ox_, LAG_TFA_ and LAG_TFA_Sulf_) in % (mol mol^-1^; n=3). The composition was determined after aldose reduction and *per*-acetylation by GC-FID/MS measurements.

| Neutral monosaccharide | LAG | LAG<sub>Ox</sub> | LAG<sub>TFA</sub> | LAG<sub>TFA_Sulf</sub> |
| --- | --- | --- | --- | --- |
| Rha | tr | - | - | - |
| Fuc | tr | tr | - | - |
| Ara | $16.0 \pm 1.2$ | $9.8 \pm 0.5$ | $4.8 \pm 0.2$ | $4.7 \pm 0.7$ |
| Xyl | tr | - | - | - |
| Man | tr | tr | tr | tr |
| Gal | $84.0 \pm 1.2$ | $90.2 \pm 0.5$ | $95.2 \pm 0.2$ | $95.3 \pm 0.7$ |
| Glc | tr | tr | tr | - |

tr: trace value < 1%

Elemental analysis of LAG_TFA_ and LAG_TFA_Sulf_ showed only traces of sulfur in LAG_TFA_, but a sulfur content of 6.7 % in LAG_TFA_Sulf_, thus revealing that chemical sulfation was successful. This was confirmed by the occurrence of typical wavenumbers 1213, 895 and 812 in FTIR (Fig. 1a). Quantification of sulfate by a photometric method confirmed successful sulfation with a content of 7.5 % ± 0.6 (w/w) in LAG_TFA_Sulf_ (Fig. 1b and 1c).

**Fig. 1.**
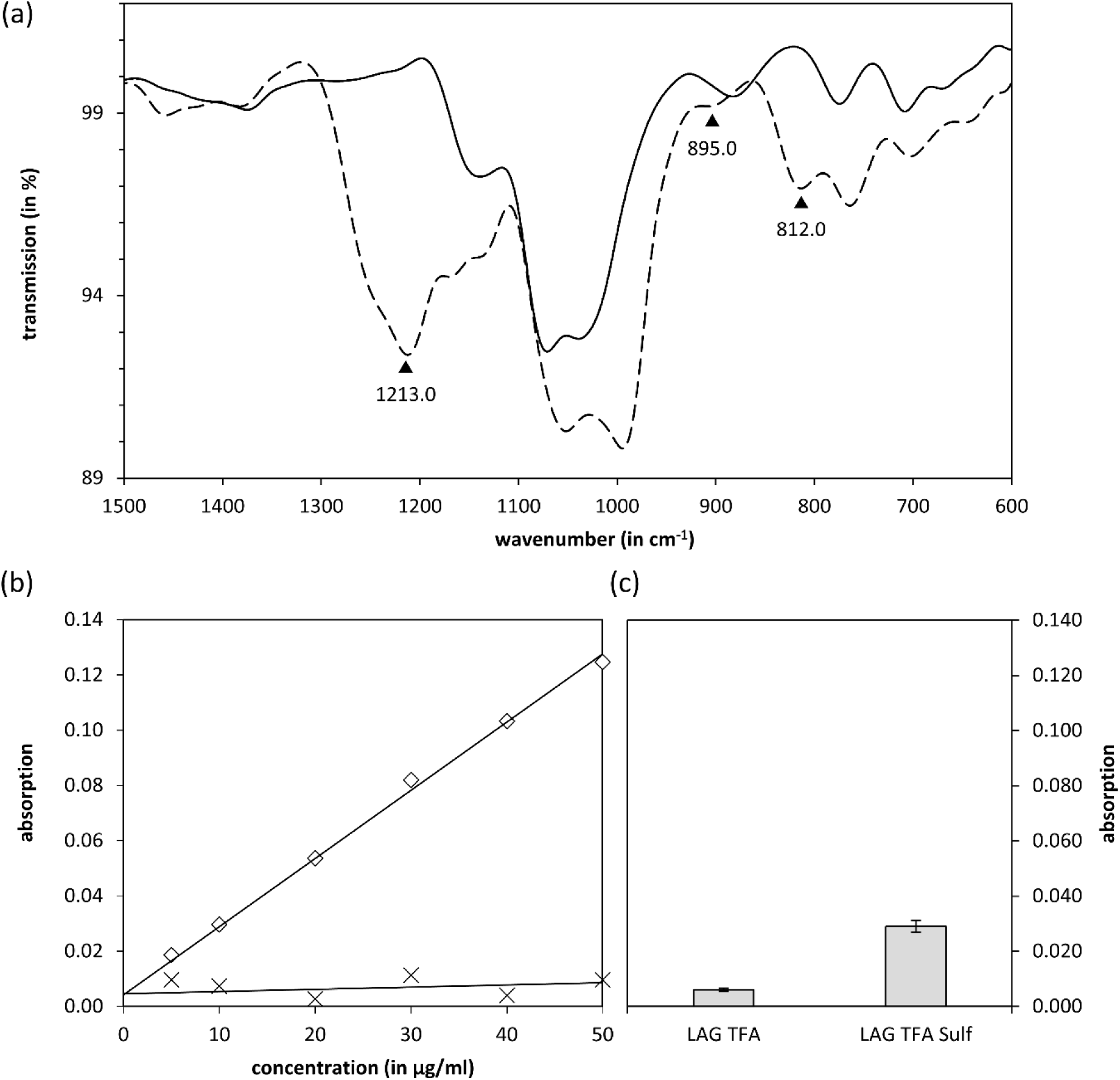
Spectrophotometric analysis of (sulfated) larch arabinogalactan samples (a) FTIR spectra of LAG_TFA_ (solid line) and LAG_TFA_Sulf_ (dashed line). (b) Calibration line of colorimetric sulfate assay using fucoidan (squares) and control line with larch arabinogalactan (crosses). (c) Measured absorption values for LAG_TFA_ and LAG_TFA_Sulf_.

### 3.2. Binding of galactans to human galectins Gal-3 and Gal-9 using BLI

The degraded AGP samples from *Echinacea* and *Zostera* (AGP_AlkOx_) as well as the TFA-treated and sulfated larch AGs (LAG_TFA_, LAG_TFA_Sulf_) were used to determine their binding affinities to human galectins Gal-3 and Gal-9. The molecular weights of the degraded AGPs from *Echinacea* and *Zostera* and the degraded LAG have been determined before and found to be 25.7 kDA for *Echinacea* AGP_AlkOx_, 29.7 kDa for *Zostera* AGP_AlkOx_ [38], 28.3 kDa for LAG_TFA_ [48] and 30.3 kDa for LAG_TFA_Sulf_. Additionally, Yariv phenylglycosides with terminal β-D-glucose (negative control; βGlcY), α-D-galactose (αGalY), β-D-galactose (βGalY), and β-D-lactose (βLacY) were synthesized ([28]; Fig. 2a) and tested as trivalent galectin inhibitors. After biotinylation, Gal-3 and Gal-9 were loaded to super-streptavidin sensors. With those, interactions with the different galactans in solution were determined by monitoring the association and dissociation curves. Sensor loading was found to be higher than 7 nm shift response for Gal-3 and higher than 3.5 for Gal-9. The steady-state method was used to calculate K_D_-values (steady-state analysis results are shown in Fig. S2-4). All samples showed concentration-dependent binding to Gal-3 with K_D_-values between 0.2 and 9.1 μmol l^-1^ (Fig. 2b and Table 4) with exception of βGlcY, which was used as negative control. Binding of degraded *Echinacea* AGP was tenfold stronger compared to the degraded *Zostera* AGP. Sulfation of LAG did not improve the binding capacity to Gal-3 (Table 4). Results were different for Gal-9 where only the degraded galactan from *Zostera* showed binding at all (Fig. 3 and Table 5).

**Fig. 2.**
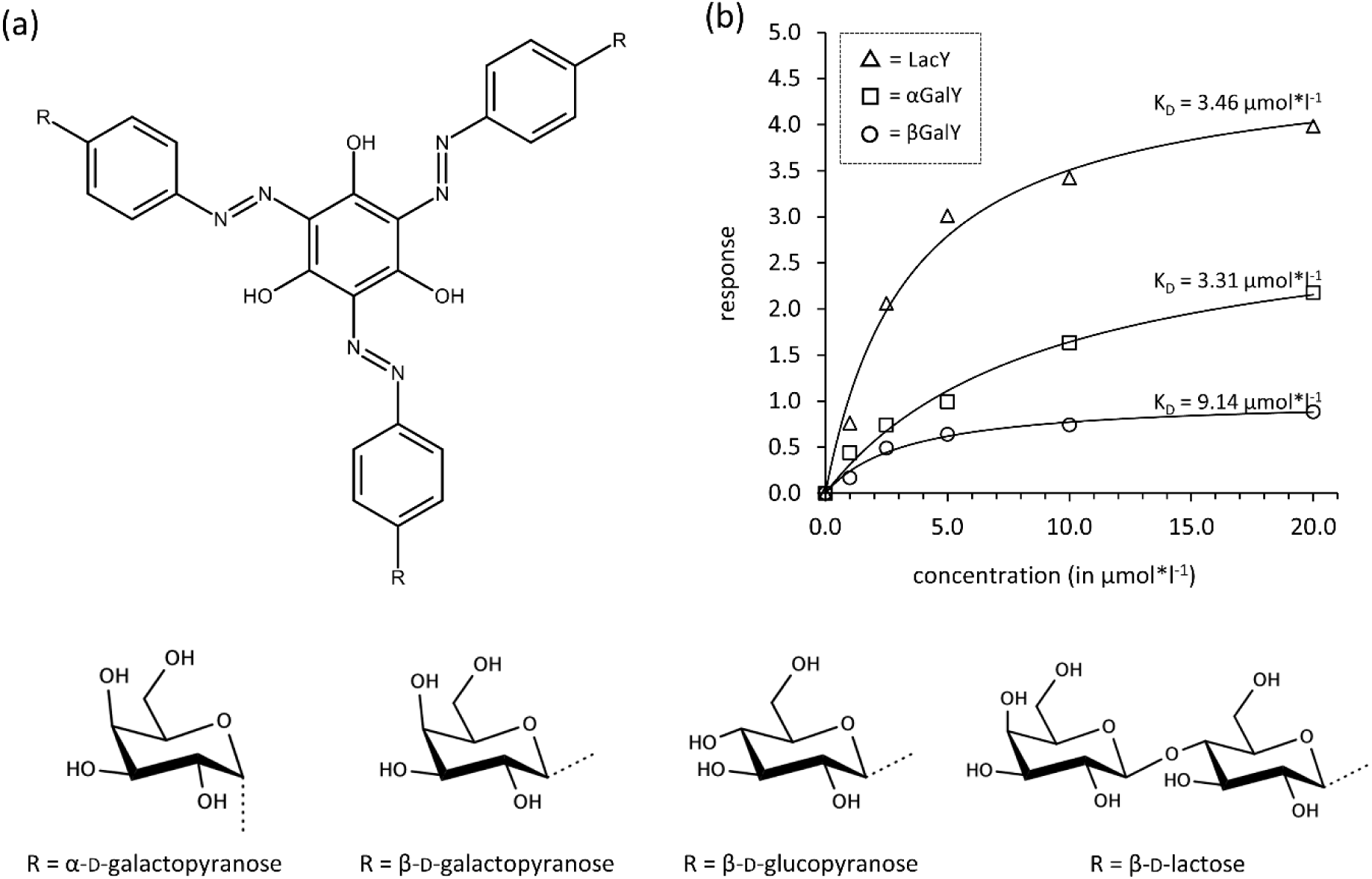
Yariv reagents as binding partners for human Gal-3. (a) Basic structure of Yariv reagents with different terminal sugar residues. (b) Steady-state analysis plot for βLacY, αGalY, βGalY. The negative control βGlcY did not bind to Gal-3.

**Fig. 3.**
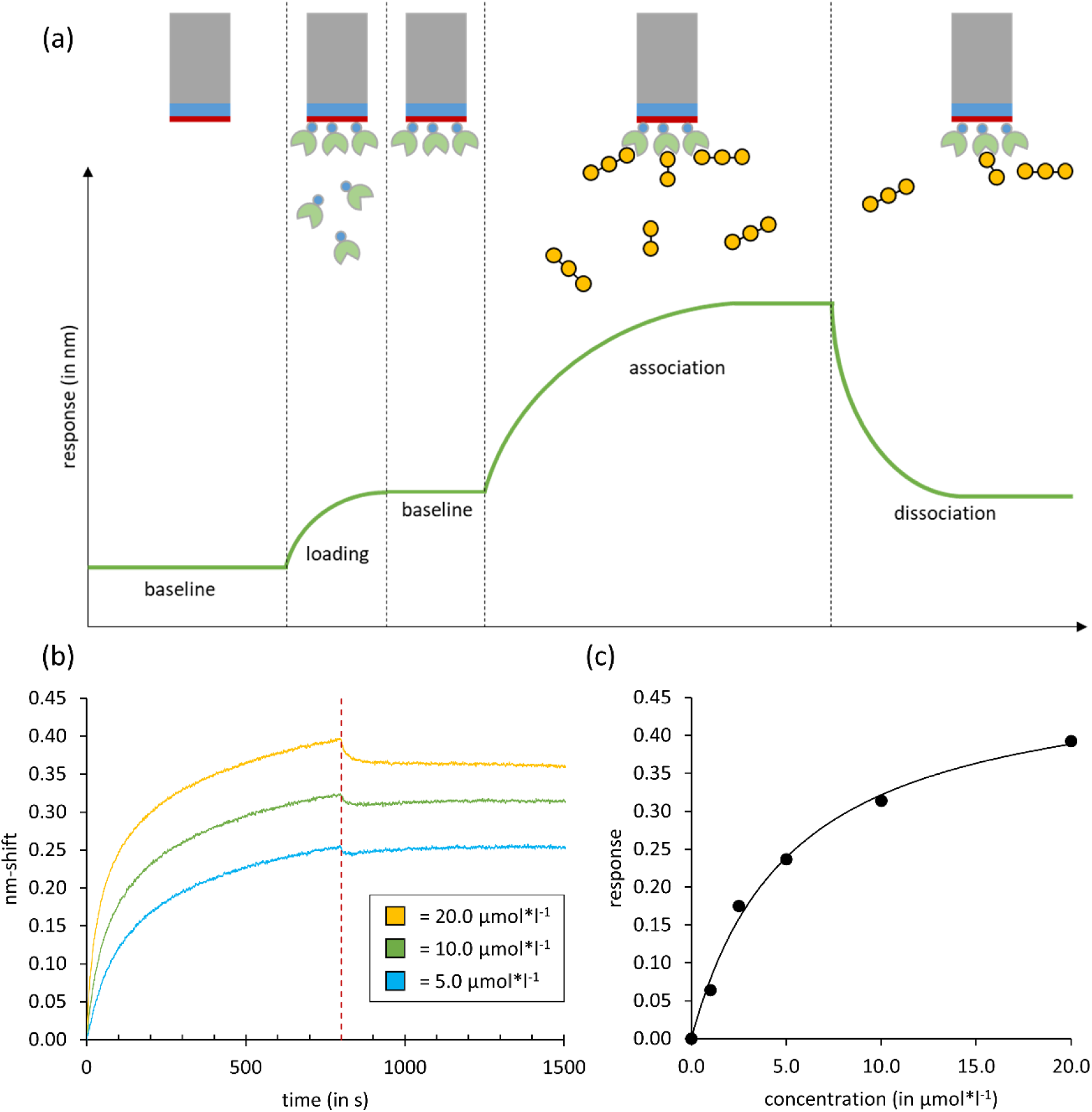
Biolayer-interferometric analysis. (a) Schematic representation of the major experimental steps during data acquisition. Sensors were loaded with tagged galectin (in green) and analysed for their binding affinities to different galactans (yellow circles). (b) Association and dissociation curves of *Zostera* galactan to Gal-9 is shown for three investigated concentrations. (c) Steady-state curve is plotted and used for K_D_ value determination.

**Table 4.** Binding of Gal-3 to different degraded plant galactans determined by BLI.

| Plant galactan | Average $K_D$-value ($\mu\text{mol l}^{-1}$) |
| --- | --- |
| *Echinacea* AGP<sub>AlkOx</sub> | 0.17 (n = 1) |
| *Zostera* AGP<sub>AlkOx</sub> | 1.82 ± 0.72 (n = 3) |
| LAG<sub>TFA</sub> | 0.40 ± 0.28 (n = 3) |
| LAG<sub>TFA_Sulf</sub> | 1.7 ± 1.27 (n = 3) |
| αGalY | 3.31 (n = 1) |
| βGalY | 9.14 ± 3.42 (n = 3) |
| βLacY | 3.46 ± 1.75 (n = 3) |
| βGlcY | no binding |

**Table 5.** Binding of Gal-9 to different degraded plant galactans determined by BLI. Average K_D_-values were calculated from the different concentrations of one experimental run.

| Plant galactan | Average $K_D$-value ($\mu\text{mol l}^{-1}$) |
| --- | --- |
| *Echinacea* AGP<sub>AlkOx</sub> | no binding |
| *Zostera* AGP<sub>AlkOx</sub> | $5.26 \pm 0.34$ (n = 3) |
| LAG<sub>TFA</sub> | no binding |
| LAG<sub>TFA_Sulf</sub> | no binding |
| αGalY | no binding |
| βGalY | no binding |
| βLacY | no binding |

### 3.3. Presence of Gal-3 and -9 in PDAC cell lines

Due to a potential of Gal-3 and -9 as therapeutic targets in PDAC, we investigated Gal-3 and -9 levels in different PDAC cell lines (Panc1 and Panc89) and their related Holo- and Paraclone variants [40,41]. Panc1 cells originate from a primary PDAC; Panc89 cells from a PDAC lymph node metastasis. Holoclones are a cancer cell variant with high amounts of cancer stem cells (CSCs), while Paraclones consist of more differentiated non-CSCs [42]. Western blot analysis revealed that Gal-3 was primarily detectable in Panc1 cell variants, with a faintly detectable signal for Panc89 cell variants, whereas Gal-9 was only present in Panc89 cell variants (Fig. 4a and 4b). Quantification of Western blot analysis (n=3) showed significantly stronger expression of Gal-3 in the Panc1 cell variants, especially in Panc1 Paraclones, where the amount of Gal-3 was more than doubled compared to the parental Panc1 cells (Fig. 4c). In contrast, Gal-9 was not detected in Panc1 cell variants, but was highly verifiable in Panc89 cell variants. Here, expression of Gal-9 was significantly lower in the Panc89 Holoclone cells compared to parental Panc89 and Panc89 Paraclone cells (Fig. 4d). Overall, these results indicate a high heterogeneity in Gal-3 and -9 expression within PDAC cell populations.

**Fig. 4.**
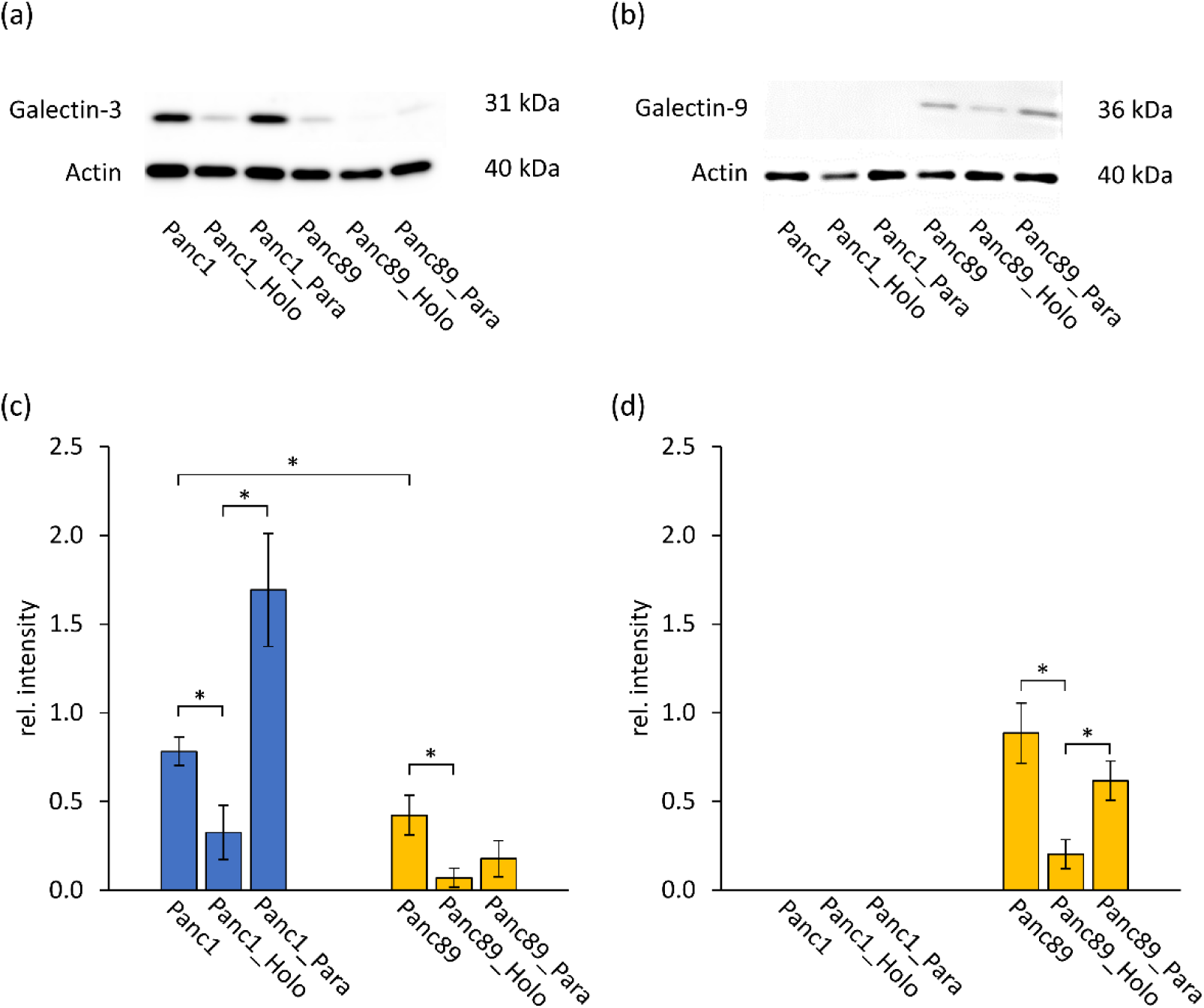
Western blot analysis of galectin-3 (Gal-3) and galectin-9 (Gal-9) on PDAC cell lines. (a), (b) Western blot of Gal-3 and Gal-9, respectively, expressed by the different investigated PDAC cell variants. (c), (d) Densitometric quantification of Gal-3 (c) and Gal-9 (d) levels relative to the reference protein actin. Significance (*p* < 0.05) is highlighted with asterisks. Global *p*-values as evaluated by one-way ANOVA were determined as *p* = 3.67e^-6^ (Gal-3) and *p* = 8.17e^-7^ (Gal-9), respectively.

## 4. Discussion

### 4.1. Plant-derived polysaccharides as potential galectin inhibitors

Especially due to their role in tumor progression, Gal-3 and Gal-9 have been identified as potential therapeutic targets. Strategies targeting galectins include small-molecule glycan inhibitors, peptides, monoclonal antibodies, truncated galectins and natural (degraded) polysaccharides [13]. In a first step, binding potentials of these inhibitors to the galectins have to be evaluated. Different methods have been established to evaluate lectin-glycan interactions, e.g. glycan microarray, isothermal calorimetry (ITC), frontal affinity chromatography, surface plasmon resonance (SPR; [49]) or biolayer interferometry (BLI; [50]). All methods have different strengths and limitations. Microarray allows high throughput, but does not provide quantitative data in terms of the binding affinity expressed by K_D_. The advantage of ITC is that both binding partners are measured in solution. Therefore, artificial immobilization does not impair affinity. This method requires substantial amounts of material [49]. Frontal affinity chromatography provides a quantitative set of K_D_s between immobilized lectins and a high number of fluorescent-labelled glycans. SPR and BLI are commonly used label-free, real-time assays with one immobilized partner, which reacts with a potential binding partner in solution. For interaction of pectin-derivatives with Gal-3, the comparability of SPR and BLI measurements has been demonstrated [51]. It has to be taken into account, that results in BLI may slightly differ dependent on usage of different sensors [51].

The galactans used in this study all belong to type-II arabinogalactans with a galactan core composed of 1,3-, 1,6- and 1,3,6-β-D-galactose. Whereas larch arabinogalactan is a pure polysaccharide [20], AGPs consist of a small protein moiety (around 10 %) which is covalently linked to several arabinogalactan polysaccharides *via O*-glycosidic linkage to hydroxyproline [21]. Native polysaccharides were degraded by mild acid to reduce the arabinose content and in case of AGPs, additionally by alkaline treatment to destroy the protein part. Compositions of degraded larch AG and degraded *Echinacea* AGP were comparable to previous investigations [38,48], whereas the residual Ara content of degraded *Zostera* AGP was higher compared to an earlier study [38]. This is probably due to the fact that starting material for larch and *Echinacea* galactans was identical to the investigations before, whereas another batch of *Zostera* rhizomes was used for isolation and degradation of AGP. Although this underlines the challenge of heterogeneity of natural plant polysaccharides *per se*, it also highlights the multivalent character of the macromolecule, which resembles the natural state of the cell surface. In further stages of drug development, a special focus should be on reproducible batch homogeneity. This could also be achieved by establishing and validating an assay to quantify drug activity to which the drug preparation can be adjusted through batch mixture.

### 4.2. Binding to Gal-3 seems to be a general feature of type I and type II galactans

In our study, all tested galactans showed comparable binding affinities to Gal-3 in a low micromolar range with strongest binding of the degraded *Echinacea* AGP. Results for this sample were comparable to previous investigations and revealed comparability of streptavidin- and super-streptavidin sensors [38]. Binding of *Zostera* was around tenfold weaker compared to degraded *Echinacea* AGP. This is explainable as the galactose content of the *Zostera* sample was around 20 % lower compared to the *Echinacea* sample (compare Tables 1 and 2).

With regard to polysaccharide galectin inhibitors, mainly modified pectins [14,15] and galactomannans [17] have been investigated up to now. In BLI, homogalacturonans which consist of a linear chain of α-(1-4)-linked D-galacturonic acid revealed lower binding affinities to Gal-3 [15] compared to rhamnogalacturonan-I [51–53]. Especially the neutral 1,4-linked galactan side chains of RG I seem to be important for binding [54–56]. Potato galactan, which also mainly consists of linear 1,4-linked galactose (type I galactan), binds to Gal-3 by its terminal, non-reducing end [57]. In our samples, structure of galactans is different with 1,3-, 1,6- and 1,3,6-linkage of galactose (type II galactan). It has been shown, that the parasite *Leishmania* possesses a poly-β-galactosyl epitope with 1,3-linked galactose which is recognized by Gal-3 and Gal-9 [25]. A strong argument that the here investigated 1,3-linked galactans might bind to galectins is the fact, that some plant β-(1,3)-galactosyltransferases possess a galectin domain [26,27].

This study provided initial insights into the influence of modification by sulfation. Although sulfation might enhance interaction with galectins [58] and binding of keratan sulfate to Gal-3 was reduced by deletion of sulfate groups [59], binding of LAG_TFA_ to Gal-3 was not improved by sulfation. This finding was unexpected but supports previous findings as one reason might be that unspecific binding of sulfate groups might block essential hydroxyl groups necessary for interaction; e.g. in galactosaminoglycans, integrity of 6-OH of βGalNAc is important for binding to galectins [60]. This reveals one limitation of our study but also of other studies working on sulfation in order to modulate activities on target proteins: homogenous sulfation of singular hydroxyl groups in a naturally-occuring polysaccharide is a challenge and knowledge on the “sulfation code” [59] would help disentangling the underlying mechanistical basics.

### 4.3. Synthetic Yariv phenylglycosides are possible Gal-3 antagonists

Yariv reagents are red-colored dyes, which are a tool to detect AGPs in plant tissues. Although the exact precipitation mechanism is still discussed, it is certain that the β-1,3-linked galactan core of AGPs is important for this interaction [61,62]. On the other hand, the structure of Yariv reagents with trivalent presentation of saccharides seems to offer a suitable structure for carbohydrate-protein interactions, in this case affinity to galectins. To date, different potential galectin inhibitors are combinations of saccharides with synthetic molecules (for review, see [12]). Some contain only one galactose residue combined with synthetic elements [63], others consist of monosaccharide-, disaccharide- or multivalent-based molecules with synthetic extensions [64].

K_D_s for binding of GalY and βLacY to Gal-3 determined in this study are in a range comparable to synthetic small molecule carbohydrate inhibitors, with exceptions revealing higher affinities, e.g. a fluoridated thio-digalactoside [65] or hexavalent clusters with LacNAc [66].

### 4.4. Binding to Gal-9 is restricted to *Zostera* galactan

Only the galactan from *Zostera* bound to Gal-9, whereas larch AG and *Echinacea* galactan as well as Yariv’s reagents showed no binding at all. This reveals that despite high structural homology of the CRDs of all galectin members, sugar-binding specificities are still present and are obviously highly important for development of selective galectin inhibitors. It has to be taken into account, that binding affinities of both CRDs (C- and N-terminal) of Gal-9 are different. Frontal affinity chromatography revealed that Gal-9N has a preference for repeated oligolactosamines and also glycolipids [67]. Investigations with synthetic galactoside and guloside derivatives revealed that galactosides selectively bound to Gal-9C, whereas the gulosides preferentially bound to Gal-9N [68]. Our study revealed a distinct binding profile of *Zostera* galactan to Gal-9 which enlightens the importance of this polysaccharide for future design of selective Gal-9 inhibitors. A future step to achieve this goal is further degradation of the galactans with subsequent chromatographic separation, followed by structural characterization and binding measurements of the resulting fractions (compare the workflow of [69]).

Special features of *Zostera* galactan are higher galactan branching represented by 1,3,6-Gal and a higher number of charged residues, namely glucuronic acids [38]. Higher charge of *Zostera* galactan is possibly the result of adaptation to the marine habitat. Branching of glycans has been identified to increase affinity to galectins [49] and Gal-9N has been reported to interact with the charged polysaccharide keratan sulfate [59]. Furthermore, the *Zostera* galactan is characterized by higher amounts of residual Ara residues, which might contribute to binding capacity, as mycobacterial arabinogalactans are recognized by Gal-9 [70]. Gal-9 was identified after isolation from tumor tissues of patients with Hodgkin’s disease [71], but its role in cancer biology is contradictory, as it seems to suppress tumor metastasis in different cases [72]. Presence of Gal-9 in cervical cancer is associated with a better overall survival [73]. In contrast, high expression of Gal-9 in tumors of metastasized PDAC patients is associated with poor survival [72]. Thus, Gal-9 might exert different roles in different cancer types either promoting or inhibiting cancer development [74]. Focusing at the monosaccharide composition, *Zostera* galactan is the only galactan investigated in this study which has a relatively high rhamnose content – a desoxyhexose being also one of the main monosaccharide components of RG-I.

Interestingly, Gal-8 (which has high similarity to Gal-9, [75]) binds to RG-I fragments [69]. L-Rhamnose is missing in mammal glycosylation patterns [77], but L-fucose is associated with several pathogenic processes [78]. The moth *Antheraea roylei* contains a lectin recognizing both, L-fucose and L-rhamnose, with stronger inhibitory effect of L-rhamnose [79]. Thus, it is not unlikely that there is cross-reactivity in substrate recognition also in human galectins. Gal-1, Gal-3, Gal-9 show substantial affinities to fucosylated structures [80,81]; e.g. Gal-1 reveals higher binding affinities to fucosylated galactose than to Galβ-1,4-GlcNAc [81]. Therefore, *Zostera* galactan might be a promising starting material to understand the relevance of desoxyhexose modification for the binding behaviour to different galectins.

### 4.5. First insights in molecular binding modes of plant saccharides to galectins

Galectins have a conserved β-sandwich-folded carbohydrate recognition domain (CRD) with a six-stranded conventional β-galactoside binding S-face and an opposing five stranded β-sheet (F-face; [16]). Gal-3 as the only chimera type galectin has an additional N-terminal tail (NT). Gal-9 belongs to the tandem repeat galectins with two distinct evolutionary conserved CRDs joined by a linker domain of variable lengths, generating isoforms [72,82]. According to [49], binding profiles of tandem-repeat galectins are comparable to the sum of the individual domains of each galectin. It has been reported that the canonical S-face (or sugar-binding face) of the CRD primarily binds to smaller β-galactosides like lactose whereas pectins and galactomannans show only poor binding to this region [18]. In contrast, the opposing noncanonical F-face can bind larger, different types of polysaccharides [15,19]. For a rhamnogalacturonan from Ginseng, multiple binding sites to Gal-3 have been proposed: one on the CRD S-face, one at the F-face and finally one within the NT [83]. Recent investigations give first hints that degradation of plant polysaccharides leads to binding of these smaller products to the S-Face of Gal-3. Partially hydrolyzed type I galactans from potato bind to the S-face with their terminal, non-reducing end; and increase of galactan chain length increased affinity [57]. Accordingly, homogalacturonan oligosaccharides interact with the S-face of Gal-3, although in an unconventional orientation [84]. Our work introduces plant arabinogalactan-proteins as new source to produce galactans of different structure capable to interact with different galectins. Their affinity values – in low micromolar ranges – are comparable to other known inhibitory substances [13] with only few substances reaching high nanomolar KDs [85]. Also natural binding partners are, with very few exceptions, in these affinity ranges [86]. Future work should aim at further degradation and elucidation of binding features of these type II galactans to galectins. In general, because of the good water solubility, absence of toxicity and multivalent display of ligands, plant derived galactans might serve as starting material to develop carbohydrate-based galectin antagonists. As search for galectin inhibitors selective for one galectin is a major challenge, binding mode of *Zostera* galactan to Gal-9 should be further investigated.

### 4.6. PDAC cell lines differ with regard to galectin expression

The analysis of protein levels of Gal-3 and -9 in PDAC cell lines revealed a high heterogeneity between the two cell lines but also among the different cell variants. While Gal-3 was predominantly expressed by Panc1 cells being characterized by a mesenchymal-like phenotype, Gal-9 was only detectable in Panc89 cells exhibiting an epithelial phenotype [40]. Heterogeneous expression of galectins in different cell types from PDAC patient biopsies were demonstrated in an innovative single-cell RNA sequencing approach [87] thus being in line with our result that Gal-9 is expressed in PDAC cell lines. Gal-3 has been already associated with Epithelial-Mesenchymal-Transition (EMT), a process that confers a motile and invasive phenotype to carcinoma cells. In colon cancer, elevated Gal-3 expression was found to be correlated with tumor aggressiveness and lowered expression of the epithelial marker E-cadherin [88]. Panc1 cells are also characterized by an EMT phenotype implying negligible E-cadherin expression. Furthermore, it was shown that Gal-3 mediates adhesion of cancer cells to liver endothelial cells [89]. Accordingly, the different migration and invasion abilities of Panc1 and Panc89 cells [40] might be related to the differing expression patterns of Gal-3 in these cells. Inhibition of Gal-3 and Gal-9 was shown to have varying, but only moderate, influence on the proliferation of PDAC cell lines ([8,29], unpublished own data). Therefore, these galectins seem to impact progression and metastasis by other processes (e.g. migration and adhesion) [89]. First therapeutic strategies targeting Gal-9 alone or together with immune checkpoint inhibitors in preclinical PDAC models revealed considerable anti-tumor effects [92,93]. There are first results showing that PDAC patients with metastasis at the time of diagnosis can be grouped into long-term (>12 months) and short-term (<12 month) survivors by using Gal-9 serum levels as novel biomarkers [8]. This could hint towards a personalized therapeutic benefit for some PDAC patients using Gal-9 specific inhibitors. In line with the immune regulatory role, the higher Gal-9 expression in Panc89 cells compared to Panc1 cells might be explained by the origin of both cell lines. While Panc1 cells are derived from the primary tumor in which the cells have presumably undergone EMT but not having spread, Panc89 cells originate from a lymph node metastasis. Thus, these cells have successfully disseminated to lymph nodes thereby encountering different environments with a plethora of immune cells requiring effective strategies surviving immune cell’s attack, e.g. upregulation of Gal-9. However, our data also reveal clear expression differences between the cell variants of either cell line being in line with the fact that a tumor comprises different phenotypic cells. Even though more PDAC cell lines could be screened, the analysis of two PDAC cell lines with different subclones derived thereof supported not only interpatient tumor heterogeneity but also intratumor heterogeneity observed in a patient tumor. The heterogeneity has to be seen as both, a limiting factor for therapeutic strategies but also as a chance to design specific inhibitors providing some patients a more personalized and effective therapy. The aim of this study was to give first insights into the potential of galactans as tumor associated inhibitors. To translate these findings into clinical application further studies beyond *in-vitro* studies have to be performed, e.g. using patient-derived multicellular tumor models or *in-vivo* studies.

## 5. Conclusions

Galectins are attractive targets in cancer therapy. In this comprehensive approach, combining biochemical and cellular analyses, we investigated the potential of a novel group of inhibitors for Gal-3 and Gal-9: chemically degraded galactose-rich molecules of plant origin. All molecules bound to Gal-3 in the micromolar ranges but affinity to Gal-9 was restricted to *Zostera* galactan. As Gal-3 and Gal-9 are involved in progression of PDAC, presence of both galectins on two PDAC cell lines and distinct cell variants thereof was evaluated. Whereas both cell lines expressed Gal-3, Gal-9 was only detected in Panc89 cells. Therefore, optimization of the *Zostera* galactan as lead structure has the potential to get closer to more personalized therapeutic options for PDAC types with overexpression of Gal-9. These findings support the view of a heterogeneous expression of galectins in PDAC cell populations, which also underline the need for further diversification of the palette of therapeutic regimes in a stratified manner.

## Abbreviations

αGalY: α-galactosyl Yariv;
AE: aqueous extract;
AG: arabinogalactan;
AGP: arabinogalactan-protein;
βGalY: β-galactosyl Yariv;
βGlcY: β-glucosyl Yariv;
βLacY: β-lactosyl Yariv;
BLI: biolayer interferometry;
CSCs: cancer stem cells;
CRD: carbohydrate recognition domain;
EMT: epithelial-mesenchymal-transition;
Gal: galectin;
ITC: isothermal calorimetry;
LAG: larch arabinogalactan;
PDAC: pancreatic ductal adenocarcinoma;
SPR: surface plasmon resonance

## Acknowledgements

A doctoral scholarship for Maximilian Thal Müller funded by the Medical Faculty of Kiel University is acknowledged.

## Author contributions: CRediT

Lukas Pfeifer: Conceptualization, data curation, formal analysis, investigation, methodology, visualization, writing-original draft, writing: review and editing

Kim-Kristine Mueller: Data curation, formal analysis, investigation, methodology, writing-review and editing

Maximilian Thal Müller: Data curation, investigation, writing-review and editing

Lisa-Marie Philipp: Data curation, formal analysis, investigation, methodology, writing-original draft, writing-review and editing

Susanne Sebens: Conceptualization, formal analysis, funding acquisition, methodology, resources, supervision, writing-original draft, writing: review and editing

Birgit Classen: Conceptualization, formal analysis, funding acquisition, methodology, project administration, resources, supervision, writing-original draft, writing: review and editing

## Appendix A. Supplementary data

Supplementary data to this article can be found online.

## References

[1] R.D. Cummings, F.-T. Liu, G.A. Rabinovich, S.R. Stowell, G.R. Vasta, Chapter 36: Galectins, in: A. Varki, R.D. Cummings, J.D. Esko, P. Stanley, G.W. Hart, M. Aebi, D. Mohnen, T. Kinoshita, N.H. Packer, J.H. Prestegard, R.L. Schnaar, P.H. Seeberger (Eds.), Essentials of Glycobiology: Galectins, 4th ed., Cold Spring Harbor (NY), 2022.

[2] L. Johannes, R. Jacob, H. Leffler, Galectins at a glance, Journal of cell science 131 (2018). 10.1242/jcs.208884.

[3] H. Verkerke, M. Dias-Baruffi, R.D. Cummings, C.M. Arthur, S.R. Stowell, Galectins: An Ancient Family of Carbohydrate Binding Proteins with Modern Functions, Methods in molecular biology (Clifton, N.J.) 2442 (2022) 1–40. 10.1007/978-1-0716-2055-7_1.

[4] L.S. Lau, N.B.B. Mohammed, C.J. Dimitroff, Decoding Strategies to Evade Immunoregulators Galectin-1, -3, and -9 and Their Ligands as Novel Therapeutics in Cancer Immunotherapy, International journal of molecular sciences 23 (2022). 10.3390/ijms232415554.

[5] M.R. Girotti, M. Salatino, T. Dalotto-Moreno, G.A. Rabinovich, Sweetening the hallmarks of cancer: Galectins as multifunctional mediators of tumor progression, The Journal of experimental medicine 217 (2020). 10.1084/jem.20182041.

[6] Z. Jiang, W. Zhang, G. Sha, D. Wang, D. Tang, Galectins Are Central Mediators of Immune Escape in Pancreatic Ductal Adenocarcinoma, Cancers 14 (2022). 10.3390/cancers14225475.

[7] D. Daley, V.R. Mani, N. Mohan, N. Akkad, A. Ochi, D.W. Heindel, K.B. Lee, C.P. Zambirinis, G.S.B. Pandian, S. Savadkar, A. Torres-Hernandez, S. Nayak, D. Wang, M. Hundeyin, B. Diskin, B. Aykut, G. Werba, R.M. Barilla, R. Rodriguez, S. Chang, L. Gardner, L.K. Mahal, B. Ueberheide, G. Miller, Dectin 1 activation on macrophages by galectin 9 promotes pancreatic carcinoma and peritumoral immune tolerance, Nature medicine 23 (2017) 556–567. 10.1038/nm.4314.

[8] A.M. Seifert, C. Reiche, M. Heiduk, A. Tannert, A.-C. Meinecke, S. Baier, J. von Renesse, C. Kahlert, M. Distler, T. Welsch, C. Reissfelder, D.E. Aust, G. Miller, J. Weitz, L. Seifert, Detection of pancreatic ductal adenocarcinoma with galectin-9 serum levels, Oncogene 39 (2020) 3102–3113. 10.1038/s41388-020-1186-7.

[9] J.-H. Chen, R.-Z. Ni, M.-B. Xiao, J.-G. Guo, J.-W. Zhou, Comparative proteomic analysis of differentially expressed proteins in human pancreatic cancer tissue, Hepatobiliary & pancreatic diseases international HBPD INT 8 (2009) 193–200.

[10] S. Song, B. Ji, V. Ramachandran, H. Wang, M. Hafley, C. Logsdon, R.S. Bresalier, Overexpressed galectin-3 in pancreatic cancer induces cell proliferation and invasion by binding Ras and activating Ras signaling, PloS one 7 (2012) e42699. 10.1371/journal.pone.0042699.

[11] L. Xie, W.-K. Ni, X.-D. Chen, M.-B. Xiao, B.-Y. Chen, S. He, C.-H. Lu, X.-Y. Li, F. Jiang, R.-Z. Ni, The expressions and clinical significances of tissue and serum galectin-3 in pancreatic carcinoma, Journal of cancer research and clinical oncology 138 (2012) 1035–1043. 10.1007/s00432-012-1178-2.

[12] D. Laaf, P. Bojarová, L. Elling, V. Křen, Galectin-Carbohydrate Interactions in Biomedicine and Biotechnology, Trends in biotechnology 37 (2019) 402–415. 10.1016/j.tibtech.2018.10.001.

[13] K.V. Mariño, A.J. Cagnoni, D.O. Croci, G.A. Rabinovich, Targeting galectin-driven regulatory circuits in cancer and fibrosis, Nature reviews. Drug discovery 22 (2023) 295–316. 10.1038/s41573-023-00636-2.

[14] E.G. Maxwell, N.J. Belshaw, K.W. Waldron, V.J. Morris, Pectin – An emerging new bioactive food polysaccharide, Trends in Food Science & Technology 24 (2012) 64–73. 10.1016/j.tifs.2011.11.002.

[15] T. Zhang, M.C. Miller, Y. Zheng, Z. Zhang, H. Xue, D. Zhao, J. Su, K.H. Mayo, Y. Zhou, G. Tai, Macromolecular assemblies of complex polysaccharides with galectin-3 and their synergistic effects on function, The Biochemical journal 474 (2017) 3849– 3868. 10.1042/BCJ20170143.

[16] L.d.F. Pedrosa, A. Raz, J.P. Fabi, The Complex Biological Effects of Pectin: Galectin-3 Targeting as Potential Human Health Improvement?, Biomolecules 12 (2022). 10.3390/biom12020289.

[17] N. Demotte, R. Bigirimana, G. Wieërs, V. Stroobant, J.-L. Squifflet, J. Carrasco, K. Thielemans, J.-F. Baurain, P. van der Smissen, P.J. Courtoy, P. van der Bruggen, A short treatment with galactomannan GM-CT-01 corrects the functions of freshly isolated human tumor-infiltrating lymphocytes, Clinical cancer research an official journal of the American Association for Cancer Research 20 (2014) 1823–1833. 10.1158/1078-0432.CCR-13-2459.

[18] J. Stegmayr, A. Lepur, B. Kahl-Knutson, M. Aguilar-Moncayo, A.A. Klyosov, R.A. Field, S. Oredsson, U.J. Nilsson, H. Leffler, Low or No Inhibitory Potency of the Canonical Galectin Carbohydrate-binding Site by Pectins and Galactomannans, The Journal of biological chemistry 291 (2016) 13318–13334. 10.1074/jbc.M116.721464.

[19] M.C. Miller, H. Ippel, D. Suylen, A.A. Klyosov, P.G. Traber, T. Hackeng, K.H. Mayo, Binding of polysaccharides to human galectin-3 at a noncanonical site in its carbohydrate recognition domain, Glycobiology 26 (2016) 88–99. 10.1093/glycob/cwv073.

[20] E.M. Göllner, H. Ichinose, S. Kaneko, W. Blaschek, B. Classen, An arabinogalactan-protein from whole grain of Avena sativa L. belongs to the wattle-blossom type of arabinogalactan-proteins, Journal of Cereal Science 53 (2011) 244–249. 10.1016/j.jcs.2011.01.004.

[21] Y. Ma, W. Zeng, A. Bacic, K. Johnson, AGPs Through Time and Space, in: J.A. Roberts (Ed.), Annual Plant Reviews online, Wiley, 2018, pp. 767–804.

[22] A.E. Clarke, R.L. Anderson, B.A. Stone, Form and function of arabinogalactans and arabinogalactan-proteins, Phytochemistry 18 (1979) 521–540. 10.1016/S0031-9422(00)84255-7.

[23] B. Classen, K. Witthohn, W. Blaschek, Characterization of an arabinogalactan-protein isolated from pressed juice of Echinacea purpurea by precipitation with the beta-glucosyl Yariv reagent, Carbohydrate research 327 (2000) 497–504. 10.1016/s0008-6215(00)00074-4.

[24] L. Pfeifer, T. Shafee, K.L. Johnson, A. Bacic, B. Classen, Arabinogalactan-proteins of Zostera marina L. contain unique glycan structures and provide insight into adaption processes to saline environments, Scientific reports 10 (2020) 8232. 10.1038/s41598-020-65135-5.

[25] I. Pelletier, T. Hashidate, T. Urashima, N. Nishi, T. Nakamura, M. Futai, Y. Arata, K. Kasai, M. Hirashima, J. Hirabayashi, S. Sato, Specific recognition of Leishmania major poly-beta-galactosyl epitopes by galectin-9: possible implication of galectin-9 in interaction between L. major and host cells, The Journal of biological chemistry 278 (2003) 22223–22230. 10.1074/jbc.M302693200.

[26] Y. Qu, J. Egelund, P.R. Gilson, F. Houghton, P.A. Gleeson, C.J. Schultz, A. Bacic, Identification of a novel group of putative Arabidopsis thaliana beta-(1,3)-galactosyltransferases, Plant molecular biology 68 (2008) 43–59. 10.1007/s11103-008-9351-3.

[27] A.M. Showalter, D. Basu, Glycosylation of arabinogalactan-proteins essential for development in Arabidopsis, Communicative & integrative biology 9 (2016) e1177687. 10.1080/19420889.2016.1177687.

[28] J. Yariv, M.M. Rapport, L. Graf, The interaction of glycosides and saccharides with antibody to the corresponding phenylazo glycosides, The Biochemical journal 85 (1962) 383–388. 10.1042/bj0850383.

[29] A. Hann, A. Gruner, Y. Chen, T.M. Gress, M. Buchholz, Comprehensive analysis of cellular galectin-3 reveals no consistent oncogenic function in pancreatic cancer cells, PloS one 6 (2011) e20859. 10.1371/journal.pone.0020859.

[30] B. Classen, S.-L. Mau, A. Bacic, The arabinogalactan-proteins from pressed juice of Echinacea purpurea belong to the hybrid class of hydroxyproline-rich glycoproteins, Planta medica 71 (2005) 59–66. 10.1055/s-2005-837752.

[31] A.B. Blakeney, P.J. Harris, R.J. Henry, B.A. Stone, A simple and rapid preparation of alditol acetates for monosaccharide analysis, Carbohydrate research 113 (1983) 291–299. 10.1016/0008-6215(83)88244-5.

[32] K.-K. Mueller, L. Pfeifer, L. Schuldt, P. Szövényi, S. de Vries, J. de Vries, K.L. Johnson, B. Classen, Fern cell walls and the evolution of arabinogalactan proteins in streptophytes, The Plant journal for cell and molecular biology 114 (2023) 875–894. 10.1111/tpj.16178.

[33] P.A. Gleeson, A.E. Clarke, Structural studies on the major component of Gladiolus style mucilage, an arabinogalactan-protein, The Biochemical journal 181 (1979) 607–621. 10.1042/bj1810607.

[34] T. Chen, B. Li, Y. Li, C. Zhao, J. Shen, H. Zhang, Catalytic synthesis and antitumor activities of sulfated polysaccharide from Gynostemma pentaphyllum Makino, Carbohydrate polymers 83 (2011) 554–560. 10.1016/j.carbpol.2010.08.024.

[35] L. Wang, X. Li, Z. Chen, Sulfated modification of the polysaccharides obtained from defatted rice bran and their antitumor activities, International journal of biological macromolecules 44 (2009) 211–214. 10.1016/j.ijbiomac.2008.12.006.

[36] J. Demšar, T. Curk, A. Erjavec, Č. Gorup, T. Hočevar, M. Milutinovič, M. Možina, M. Polajnar, M. Toplak, A. Starič, M. Štajdohar, L. Umek, L. Žagar, J. Žbontar, M. Žitnik, B. Zupan, Orange: Data Mining Toolbox in Python, Journal of Machine Learning Research 14 (2013) 2349–2353.

[37] R.W. Farndale, D.J. Buttle, A.J. Barrett, Improved quantitation and discrimination of sulphated glycosaminoglycans by use of dimethylmethylene blue, Biochimica et biophysica acta 883 (1986) 173–177. 10.1016/0304-4165(86)90306-5.

[38] L. Pfeifer, A. Baumann, L.M. Petersen, B. Höger, E. Beitz, B. Classen, Degraded Arabinogalactans and Their Binding Properties to Cancer-Associated Human Galectins, International journal of molecular sciences 22 (2021). 10.3390/ijms22084058.

[39] SARTORIUS Technical Note, Biotinylation of Proteins for Immobilization Onto Streptavidin Biosensors.

[40] L.-M. Philipp, U.-U. Yesilyurt, A. Surrow, A. Künstner, A.-S. Mehdorn, C. Hauser, J.-P. Gundlach, O. Will, P. Hoffmann, L. Stahmer, S. Franzenburg, H. Knaack, U. Schumacher, H. Busch, S. Sebens, Epithelial and Mesenchymal-like Pancreatic Cancer Cells Exhibit Different Stem Cell Phenotypes Associated with Different Metastatic Propensities, Cancers 16 (2024). 10.3390/cancers16040686.

[41] H. Knaack, L. Lenk, L.-M. Philipp, L. Miarka, S. Rahn, F. Viol, C. Hauser, J.-H. Egberts, J.-P. Gundlach, O. Will, S. Tiwari, W. Mikulits, U. Schumacher, J.G. Hengstler, S. Sebens, Liver metastasis of pancreatic cancer: the hepatic microenvironment impacts differentiation and self-renewal capacity of pancreatic ductal epithelial cells, Oncotarget 9 (2018) 31771–31786. 10.18632/oncotarget.25884.

[42] C.M. Beaver, A. Ahmed, J.R. Masters, Clonogenicity: holoclones and meroclones contain stem cells, PloS one 9 (2014) e89834. 10.1371/journal.pone.0089834.

[43] C.A. Schneider, W.S. Rasband, K.W. Eliceiri, NIH Image to ImageJ: 25 years of image analysis, Nature methods 9 (2012) 671–675. 10.1038/nmeth.2089.

[44] S.S. Shapiro, M.B. Wilk, An analysis of variance test for normality (complete samples), Biometrika 52 (1965) 591–611. 10.1093/biomet/52.3-4.591.

[45] A. Kassambara, rstatix: Pipe-Friendly Framework for Basic Statistical Tests, 2019, accessed 28th of November, 2024.

[46] Student, The Probable Error of a Mean, Biometrika 6 (1908) 1. 10.2307/2331554.

[47] S. Holm, A Simple Sequentially Rejective Multiple Test Procedure, Scandinavian Journal of Statistics 6 (1979) 65–70.

[48] S. André, B. Classen, H.-J. Gabius, Studies on Unprocessed and Acid-Treated Arabinogalactan from Larch as an Inhibitor of Glycan Binding of a Plant Toxin and Biomedically Relevant Human Lectins, Planta medica 81 (2015) 1146–1153. 10.1055/s-0035-1546113.

[49] J. Iwaki, J. Hirabayashi, Carbohydrate-Binding Specificity of Human Galectins: An Overview by Frontal Affinity Chromatography, Trends in Glycoscience and Glycotechnology 30 (2018) SE137–SE153. 10.4052/tigg.1728.1SE.

[50] S. Murali, R.R. Rustandi, X. Zheng, A. Payne, L. Shang, Applications of Surface Plasmon Resonance and Biolayer Interferometry for Virus-Ligand Binding, Viruses 14 (2022). 10.3390/v14040717.

[51] T. Zhang, Y. Zheng, D. Zhao, J. Yan, C. Sun, Y. Zhou, G. Tai, Multiple approaches to assess pectin binding to galectin-3, International journal of biological macromolecules 91 (2016) 994–1001. 10.1016/j.ijbiomac.2016.06.058.

[52] J. Su, Y. Wang, Y. Si, J. Gao, C. Song, L. Cui, R. Wu, G. Tai, Y. Zhou, Galectin-13, a different prototype galectin, does not bind β-galacto-sides and forms dimers via intermolecular disulfide bridges between Cys-136 and Cys-138, Scientific reports 8 (2018) 980. 10.1038/s41598-018-19465-0.

[53] L. Cui, J. Wang, R. Huang, Y. Tan, F. Zhang, Y. Zhou, L. Sun, Analysis of pectin from Panax ginseng flower buds and their binding activities to galectin-3, International journal of biological macromolecules 128 (2019) 459–467. 10.1016/j.ijbiomac.2019.01.129.

[54] X. Gao, Y. Zhi, L. Sun, X. Peng, T. Zhang, H. Xue, G. Tai, Y. Zhou, The inhibitory effects of a rhamnogalacturonan I (RG-I) domain from ginseng pectin on galectin-3 and its structure-activity relationship, The Journal of biological chemistry 288 (2013) 33953–33965. 10.1074/jbc.M113.482315.

[55] A.P. Gunning, R.J.M. Bongaerts, V.J. Morris, Recognition of galactan components of pectin by galectin-3, FASEB journal official publication of the Federation of American Societies for Experimental Biology 23 (2009) 415–424. 10.1096/fj.08-106617.

[56] X. Gao, Y. Zhi, T. Zhang, H. Xue, X. Wang, A.D. Foday, G. Tai, Y. Zhou, Analysis of the neutral polysaccharide fraction of MCP and its inhibitory activity on galectin-3, Glycoconjugate journal 29 (2012) 159–165. 10.1007/s10719-012-9382-5.

[57] M.C. Miller, Y. Zheng, Y. Zhou, G. Tai, K.H. Mayo, Galectin-3 binds selectively to the terminal, non-reducing end of β(1→4)-galactans, with overall affinity increasing with chain length, Glycobiology 29 (2019) 74–84. 10.1093/glycob/cwy085.

[58] H. Ideo, A. Seko, T. Ohkura, K.L. Matta, K. Yamashita, High-affinity binding of recombinant human galectin-4 to SO(3)(-)--3Galbeta1--3GalNAc pyranoside, Glycobiology 12 (2002) 199–208. 10.1093/glycob/12.3.199.

[59] M.C. Miller, C. Cai, K. Wichapong, S. Bhaduri, N.L.B. Pohl, R.J. Linhardt, H.-J. Gabius, K.H. Mayo, Structural insight into the binding of human galectins to corneal keratan sulfate, its desulfated form and related saccharides, Scientific reports 10 (2020) 15708. 10.1038/s41598-020-72645-9.

[60] J. Iwaki, T. Minamisawa, H. Tateno, J. Kominami, K. Suzuki, N. Nishi, T. Nakamura, J. Hirabayashi, Desulfated galactosaminoglycans are potential ligands for galectins: evidence from frontal affinity chromatography, Biochemical and biophysical research communications 373 (2008) 206–212. 10.1016/j.bbrc.2008.05.190.

[61] K. Kitazawa, T. Tryfona, Y. Yoshimi, Y. Hayashi, S. Kawauchi, L. Antonov, H. Tanaka, T. Takahashi, S. Kaneko, P. Dupree, Y. Tsumuraya, T. Kotake, β-galactosyl Yariv reagent binds to the β-1,3-galactan of arabinogalactan proteins, Plant physiology 161 (2013) 1117–1126. 10.1104/pp.112.211722.

[62] B.S. Paulsen, D.J. Craik, D.E. Dunstan, B.A. Stone, A. Bacic, The Yariv reagent: behaviour in different solvents and interaction with a gum arabic arabinogalactan-protein, Carbohydrate polymers 106 (2014) 460–468. 10.1016/j.carbpol.2014.01.009.

[63] F.R. Zetterberg, K. Peterson, R.E. Johnsson, T. Brimert, M. Håkansson, D.T. Logan, H. Leffler, U.J. Nilsson, Monosaccharide Derivatives with Low-Nanomolar Lectin Affinity and High Selectivity Based on Combined Fluorine-Amide, Phenyl-Arginine, Sulfur-π, and Halogen Bond Interactions, ChemMedChem 13 (2018) 133–137. 10.1002/cmdc.201700744.

[64] V.L. Campo, M.F. Marchiori, L.C. Rodrigues, M. Dias-Baruffi, Synthetic glycoconjugates inhibitors of tumor-related galectin-3: an update, Glycoconjugate journal 33 (2016) 853–876. 10.1007/s10719-016-9721-z.

[65] T.-J. Hsieh, H.-Y. Lin, Z. Tu, T.-C. Lin, S.-C. Wu, Y.-Y. Tseng, F.-T. Liu, S.-T.D. Hsu, C.-H. Lin, Dual thio-digalactoside-binding modes of human galectins as the structural basis for the design of potent and selective inhibitors, Scientific reports 6 (2016) 29457. 10.1038/srep29457.

[66] M. Hovorková, J. Červený, L. Bumba, H. Pelantová, J. Cvačka, V. Křen, O. Renaudet, D. Goyard, P. Bojarová, Advanced high-affinity glycoconjugate ligands of galectins, Bioorganic chemistry 131 (2023) 106279. 10.1016/j.bioorg.2022.106279.

[67] J. Hirabayashi, T. Hashidate, Y. Arata, N. Nishi, T. Nakamura, M. Hirashima, T. Urashima, T. Oka, M. Futai, W.E.G. Muller, F. Yagi, K. Kasai, Oligosaccharide specificity of galectins: a search by frontal affinity chromatography, Biochimica et biophysica acta 1572 (2002) 232–254. 10.1016/s0304-4165(02)00311-2.

[68] M. Mahanti, K.B. Pal, A.P. Sundin, H. Leffler, U.J. Nilsson, Epimers Switch Galectin-9 Domain Selectivity: 3N-Aryl Galactosides Bind the C-Terminal and Gulosides Bind the N-Terminal, ACS medicinal chemistry letters 11 (2020) 34–39. 10.1021/acsmedchemlett.9b00396.

[69] Y. Zheng, Y. Si, X. Xu, H. Gu, Z. He, Z. Zhao, Z. Feng, J. Su, K.H. Mayo, Y. Zhou, G. Tai, Ginseng-derived type I rhamnogalacturonan polysaccharide binds to galectin-8 and antagonizes its function, Journal of ginseng research 48 (2024) 202–210. 10.1016/j.jgr.2023.11.007.

[70] X. Wu, Y. Wu, R. Zheng, F. Tang, L. Qin, D. Lai, L. Zhang, L. Chen, B. Yan, H. Yang, Y. Wang, F. Li, J. Zhang, F. Wang, L. Wang, Y. Cao, M. Ma, Z. Liu, J. Chen, X. Huang, J. Wang, R. Jin, P. Wang, Q. Sun, W. Sha, L. Lyu, P. Moura-Alves, A. Dorhoi, G. Pei, P. Zhang, J. Chen, S. Gao, F. Randow, G. Zeng, C. Chen, X.-S. Ye, S.H.E. Kaufmann, H. Liu, B. Ge, Sensing of mycobacterial arabinogalactan by galectin-9 exacerbates mycobacterial infection, EMBO reports 22 (2021) e51678. 10.15252/embr.202051678.

[71] O. Türeci, H. Schmitt, N. Fadle, M. Pfreundschuh, U. Sahin, Molecular definition of a novel human galectin which is immunogenic in patients with Hodgkin’s disease, The Journal of biological chemistry 272 (1997) 6416–6422. 10.1074/jbc.272.10.6416.

[72] P. Moar, R. Tandon, Galectin-9 as a biomarker of disease severity, Cellular immunology 361 (2021) 104287. 10.1016/j.cellimm.2021.104287.

[73] S. Beyer, M. Wehrmann, S. Meister, T.M. Kolben, F. Trillsch, A. Burges, B. Czogalla, E. Schmoeckel, S. Mahner, U. Jeschke, T. Kolben, Galectin-8 and -9 as prognostic factors for cervical cancer, Archives of gynecology and obstetrics 306 (2022) 1211– 1220. 10.1007/s00404-022-06449-9.

[74] C.F. Rodrigues, F.A. Santos, L.A.A. Amorim, A.L.C. Da Silva, L.G.A. Marques, B.A.M. Rocha, Galectin-9 is a target for the treatment of cancer: A patent review, International journal of biological macromolecules 254 (2024) 127768. 10.1016/j.ijbiomac.2023.127768.

[75] D. Houzelstein, I.R. Gonçalves, A.J. Fadden, S.S. Sidhu, D.N.W. Cooper, K. Drickamer, H. Leffler, F. Poirier, Phylogenetic analysis of the vertebrate galectin family, Molecular biology and evolution 21 (2004) 1177–1187. 10.1093/molbev/msh082.

[76] E. Rodriguez, K. Boelaars, K. Brown, K. Madunić, T. van Ee, F. Dijk, J. Verheij, R.J.E. Li, S.T.T. Schetters, L.L. Meijer, T.Y.S. Le Large, E. Driehuis, H. Clevers, S.C.M. Bruijns, T. O’Toole, S.J. van Vliet, M.F. Bijlsma, M. Wuhrer, G. Kazemier, E. Giovannetti, J.J. Garcia-Vallejo, Y. van Kooyk, Analysis of the glyco-code in pancreatic ductal adenocarcinoma identifies glycan-mediated immune regulatory circuits, Communications biology 5 (2022) 41. 10.1038/s42003-021-02934-0.

[77] S. Li, F. Chen, Y. Li, L. Wang, H. Li, G. Gu, E. Li, Rhamnose-Containing Compounds: Biosynthesis and Applications, Molecules (Basel, Switzerland) 27 (2022). 10.3390/molecules27165315.

[78] Z. Tu, Y.-N. Lin, C.-H. Lin, Development of fucosyltransferase and fucosidase inhibitors, Chemical Society reviews 42 (2013) 4459–4475. 10.1039/C3CS60056D.

[79] H.K. Devi, S.K. Devi, H. Rully, S.J. Singh, W.S. Singh, H. Thongam, L.R. Singh, Purification and Characterization of a Novel Rhamnose/Fucose-Specific Lectin from the Hemolymph of Oak Tasar (Antheraea proylei J.) Silkworm, Protein and peptide letters 27 (2020) 649–657. 10.2174/0929866527666200129155343.

[80] H. Makyio, T. Takeuchi, M. Tamura, K. Nishiyama, H. Takahashi, H. Natsugari, Y. Arata, K. Kasai, Y. Yamada, S. Wakatsuki, R. Kato, Structural basis of preferential binding of fucose-containing saccharide by the Caenorhabditis elegans galectin LEC-6, Glycobiology 23 (2013) 797–805. 10.1093/glycob/cwt017.

[81] T. Takeuchi, M. Tamura, K. Nishiyama, J. Iwaki, J. Hirabayashi, H. Takahashi, H. Natsugari, Y. Arata, K. Kasai, Mammalian galectins bind galactoseβ1-4fucose disaccharide, a unique structural component of protostomial N-type glycoproteins, Biochemical and biophysical research communications 436 (2013) 509–513. 10.1016/j.bbrc.2013.05.135.

[82] S. John, R. Mishra, Galectin-9: From cell biology to complex disease dynamics, Journal of biosciences 41 (2016) 507–534. 10.1007/s12038-016-9616-y.

[83] M.C. Miller, Y. Zheng, J. Yan, Y. Zhou, G. Tai, K.H. Mayo, Novel polysaccharide binding to the N-terminal tail of galectin-3 is likely modulated by proline isomerization, Glycobiology 27 (2017) 1038–1051. 10.1093/glycob/cwx071.

[84] Y. Zheng, J. Su, M.C. Miller, J. Geng, X. Xu, T. Zhang, M. Mayzel, Y. Zhou, K.H. Mayo, G. Tai, Topsy-turvy binding of negatively charged homogalacturonan oligosaccharides to galectin-3, Glycobiology 31 (2021) 341–350. 10.1093/glycob/cwaa080.

[85] J. Choutka, J. Kaminský, E. Wang, K. Parkan, R. Pohl, End-Point Affinity Estimation of Galectin Ligands by Classical and Semiempirical Quantum Mechanical Potentials, Journal of chemical information and modeling. 10.1021/acs.jcim.4c01659.

[86] M.F. Troncoso, M.T. Elola, A.G. Blidner, L. Sarrias, M.V. Espelt, G.A. Rabinovich, The universe of galectin-binding partners and their functions in health and disease, The Journal of biological chemistry 299 (2023) 105400. 10.1016/j.jbc.2023.105400.

[87] O. Abudu, D. Nguyen, I. Millward, J.E. Manning, M. Wahid, A. Lightfoot, F. Marcon, R. Merard, S. Margielewska-Davies, K. Roberts, R. Brown, S. Powell-Brett, S.M. Nicol, F. Zayou, W.D. Croft, H. Pearce, P. Moss, A.J. Iqbal, H.M. McGettrick, Interplay in galectin expression predicts patient outcomes in a spatially restricted manner in PDAC, Biomedicine & pharmacotherapy = Biomedecine & pharmacotherapie 172 (2024) 116283. 10.1016/j.biopha.2024.116283.

[88] Z. Huang, Z. Ai, N. Li, H. Xi, X. Gao, F. Wang, X. Tan, H. Liu, Over expression of galectin-3 associates with short-term poor prognosis in stage II colon cancer, Cancer biomarkers section A of Disease markers 17 (2016) 445–455. 10.3233/CBM-160661.

[89] M. Schöll-Naderer, O. Helm, J. Spencker, L. Pfeifer, T. Rätsch, S. Sebens, B. Classen, Plant-derived saccharides and their inhibitory potential on metastasis associated cellular processes of pancreatic ductal adenocarcinoma cells, Carbohydrate research 490 (2020) 107903. 10.1016/j.carres.2019.107903.

[90] C. Yıldırım, Galectin-9, a pro-survival factor inducing immunosuppression, leukemic cell transformation and expansion, Molecular biology reports 51 (2024) 571. 10.1007/s11033-024-09563-w.

[91] N.-I. Kapetanakis, P. Busson, Galectins as pivotal components in oncogenesis and immune exclusion in human malignancies, Frontiers in immunology 14 (2023) 1145268. 10.3389/fimmu.2023.1145268.

[92] E. Li, J. Xu, Q. Chen, X. Zhang, X. Xu, T. Liang, Galectin-9 and PD-L1 antibody blockade combination therapy inhibits tumour progression in pancreatic cancer, Immunotherapy 15 (2023) 135–147. 10.2217/imt-2021-0075.

[93] A. Quilbe, R. Mustapha, B. Duchêne, A. Kumar, E. Werkmeister, E. Leteurtre, O. Moralès, N. Jonckheere, I. van Seuningen, N. Delhem, A novel anti-galectin-9 immunotherapy limits the early progression of pancreatic neoplastic lesions in transgenic mice, Frontiers in immunology 14 (2023) 1267279. 10.3389/fimmu.2023.1267279.

