## Supplementary Material for "Synthetic and plant-derived multivalent galactans as modulators of cancer-associated galectins-3 and -9"

**Table S1:** Used galectin samples

| Protein | Origin | Information |
|---|---|---|
| Galectin-3, human, unconjugated | produced by Biorbyt Ltd., Cambridge, UK | recombinant protein, produced in *E. coli* |
| Galectin-9, human unconjugated | produced by Biorbyt Ltd., Cambridge, UK | recombinant protein (amino acids Ala 2 – Thr 323), produced in HEK293 cells |

**Table S2:** Used cell culture media and buffers

| Cell culture media and buffer | Composition |
|---|---|
| Panc-medium | RPMI 1640 (Biochrom GmbH, Berlin, Germany) supplemented with 10% fetal calf serum, 1% L-glutamine, and 1% sodium pyruvate |
| RIPA buffer | 50 mmol $l^{-1}$ TRIS-HCl, 150 mmol $l^{-1}$ NaCl, octylphenoxypolyethoxyethanol (IGEPAL CA-630, (Sigma-Aldrich Corp., St. Louis, MO, USA), 0.5% (w/V) sodium deoxycholate, 0.2% (w/V) sodium dodecyl sulfate |
| cell lysis buffer | RIPA buffer supplemented with EDTA-free protease inhibitor cocktail (cOmplete ULTRA tablets, Roche Diagnostics International AG, Rotkreuz, Switzerland) and phosphatase inhibitor cocktail (PhosSTOP tablets, Roche Diagnostics International AG) |
| loading dye | 8% (w/V) sodium dodecyl sulfate, 200 mmol $l^{-1}$ dithiothreitol, 40% (V/V) glycerol, a spatula tip of bromophenol blue in TRIS-HCl (250 mmol $l^{-1}$) |
| TBS (TRIS-buffered saline, 10x) | 24.4 g TRIS base, 80g NaCl, adjusted to pH 7.6 with HCl in 1l ultrapure water |
| TBST (TBS + Tween 20) | 0.1% (m/V) Tween 20 in TBS (1x) |
| PBS (phosphate-buffered saline) | 137 mmol $l^{-1}$ NaCl, 2.7 mmol $l^{-1}$ KCl, 8.1 mmol $l^{-1}$ $Na_2HPO_4$, 1.8 mmol $l^{-1}$ $KH_2PO_4$, adjusted to pH 7.4 |

**Table S3:** Methodological Steps for Biolayer Interferometry measurement

| Assay step | Assay time | Used solution |
|---|---|---|
| Wash | 600 sec | PBS (pH 7.4) |
| Wash 2 | 600 sec | PBS (pH 7.4) |
| Baseline | 1200 sec | PBS (pH 7.4) |
| Loading | 2000 sec | Galectin solution (10 µg $ml^{-1}$ Gal-3 or Gal-9) in PBS (pH 7.4) |

| | | |
|---|---|---|
| Blocking | 300 sec | blocking solution containing biocytin (10 µg $ml^{-1}$) in PBS (pH 7.4) |
| Wash 3 | 600 sec | PBS (pH 7.4) |
| Baseline 2 | 2000 sec | PBS (pH 7.4) |
| Association | 800 sec | Samples in PBS (pH 7.4) |
| Dissociation | 1200 sec | PBS (pH 7.4) |

**Table S4:** Used cell lines

| Cell line | Origin | Information |
|---|---|---|
| Panc1 | Purchased from ATCC, Manassas, VA, USA | originating from a primary PDAC |
| Panc1_Holo | Derived from single-cell cloning in the lab of Prof. S. Sebens, Kiel | Detailed description of the isolation in the original publication [1] |
| Panc1_Para | Derived from single-cell cloning in the lab of Prof. S. Sebens, Kiel | Detailed description of the isolation in the original publication [1] |
| Panc89 | Kindly provided by Prof. T. Okabe, University of Tokyo, Tokyo, Japan | originating from a PDAC lymph node metastasis |
| Panc89_Holo | Derived from single-cell cloning in the lab of Prof. S. Sebens, Kiel | Detailed description of the isolation in the original publication [2] |
| Panc89_Para | Derived from single-cell cloning in the lab of Prof. S. Sebens, Kiel | Detailed description of the isolation in the original publication [2] |

**Table S5:** Antibodies for Western Blot analysis

| Antibody | Information | Manufacturer |
|---|---|---|
| Primary antibody against galectin-9 | polyclonal antibody (IgG), produced in rabbit. Immunogen was a synthesized peptide derived from human LGALS9, corresponding to amino acid residues F60-F110 of human galectin-9. | Product number: PA5-115266 ThermoFisher Scientific GmbH, Darmstadt, Germany |
| Primary antibody against galectin-3 | monoclonal antibody ($IgG_1$ κ), produced in mouse. Epitope mapping was within the first 18 amino acids of human galectin-3. | Product number: sc-32790 Santa Cruz Biotechnology, Inc., Dallas, TX, USA |

| | | |
|---|---|---|
| Primary antibody against β-Actin | monoclonal reference antibody (mouse-$IgG_1$-isoype), clone AC-15, ascites fluid. | Product number: A5441<br>Sigma-Aldrich Chemie GmbH<br>Taufkirchen, Germany |
| Anti-rabbit secondary antibody | polyclonal antibody, produced in goat, horseradish peroxidase (HRP)-conjugated. Directed against IgG. | Product number: 7074<br>Cell Signaling Technology Europe, B.V., Leiden, Netherlands |
| Anti-mouse secondary antibody | polyclonal antibody, produced in horse, horseradish peroxidase (HRP)-conjugated. Directed against IgG. | Product number: 7076<br>Cell Signaling Technology Europe, B.V., Leiden, Netherlands |

**Table S6:** Results of the Shapiro-Wilk evaluation of the measured protein levels

| galectin | cell variant | test statistic *W* | *p*-value | normal distribution |
|---|---|---|---|---|
| Gal-3 | Panc1 | 0.855 | 0.253 | yes, null hypothesis cannot be rejected (*p*-value > 0.05) |
| Gal-3 | Panc1_Holo | 0.861 | 0.271 | yes, null hypothesis cannot be rejected (*p*-value > 0.05) |
| Gal-3 | Panc1_Para | 0.954 | 0.587 | yes, null hypothesis cannot be rejected (*p*-value > 0.05) |
| Gal-3 | Panc89 | 0.906 | 0.406 | yes, null hypothesis cannot be rejected (*p*-value > 0.05) |
| Gal-3 | Panc89_Holo | 0.986 | 0.774 | yes, null hypothesis cannot be rejected (*p*-value > 0.05) |
| Gal-3 | Panc89_Para | 0.945 | 0.548 | yes, null hypothesis cannot be rejected (*p*-value > 0.05) |
| Gal-9 | Panc1 | - | - | yes, null hypothesis cannot be rejected (*p*-value > 0.05) |
| Gal-9 | Panc1_Holo | - | - | yes, null hypothesis cannot be rejected (*p*-value > 0.05) |
| Gal-9 | Panc1_Para | - | - | yes, null hypothesis cannot be rejected (*p*-value > 0.05) |
| Gal-9 | Panc89 | 0.980 | 0.727 | yes, null hypothesis cannot be rejected (*p*-value > 0.05) |
| Gal-9 | Panc89_Holo | 0.863 | 0.276 | yes, null hypothesis cannot be rejected (*p*-value > 0.05) |
| Gal-9 | Panc89_Para | 0.987 | 0.785 | yes, null hypothesis cannot be rejected (*p*-value > 0.05) |

**Table S7:** Results of the ANOVA tests for galectin-3 and galectin-9

| effect | DF effect | DF error | F-value | *p*-value | significance | Generalized Eta-Squared measure of effect size |
|---|---|---|---|---|---|---|
| Gal-3 | 5 | 12 | 24.289 | 3.67e-06 | *** | 0.919 |
| Gal-9 | 5 | 12 | 35.904 | 8.17e-07 | *** | 0.937 |

* *p*-value < 0.05, ** *p*-value < 0.01, *** *p*-value < 0.001, ns: *p*-value > 0.05

**Table S8:** Statistical results of the pairwise comparison (t-test) of the galectin-3 protein levels

| Group 1 | Group 2 | p-value | adjusted p-value | formatted p-value | significance |
|---|---|---|---|---|---|
| Panc1 | Panc1_Holo | 0.033 | 0.240 | 0.033 | * |
| Panc1 | Panc1_Para | 0.049 | 0.290 | 0.049 | * |
| Panc1 | Panc89 | 0.025 | 0.230 | 0.025 | * |
| Panc1 | Panc89_Holo | 0.001 | 0.014 | 0.001 | *** |
| Panc1 | Panc89_Para | 0.003 | 0.047 | 0.003 | ** |
| Panc1_Holo | Panc1_Para | 0.013 | 0.170 | 0.013 | * |
| Panc1_Holo | Panc89 | 0.515 | 0.830 | 0.515 | ns |
| Panc1_Holo | Panc89_Holo | 0.129 | 0.520 | 0.129 | ns |
| Panc1_Holo | Panc89_Para | 0.326 | 0.830 | 0.326 | ns |
| Panc1_Para | Panc89 | 0.021 | 0.210 | 0.021 | * |
| Panc1_Para | Panc89_Holo | 0.017 | 0.180 | 0.017 | * |
| Panc1_Para | Panc89_Para | 0.014 | 0.170 | 0.014 | * |
| Panc89 | Panc89_Holo | 0.030 | 0.240 | 0.030 | * |
| Panc89 | Panc89_Para | 0.085 | 0.430 | 0.085 | ns |
| Panc89_Holo | Panc89_Para | 0.278 | 0.830 | 0.278 | ns |

* *p*-value < 0.05, ** *p*-value < 0.01, *** *p*-value < 0.001, ns: *p*-value > 0.05

**Table S9:** Statistical results of the pairwise comparison (t-test) of the galectin-9 protein levels

| Group 1 | Group 2 | p-value | adjusted p-value | formatted p-value | significance |
|---|---|---|---|---|---|
| Panc1 | Panc89 | 0.018 | 0.190 | 0.018 | * |
| Panc1 | Panc89_Holo | 0.073 | 0.290 | 0.073 | ns |
| Panc1 | Panc89_Para | 0.016 | 0.190 | 0.016 | * |
| Panc1_Holo | Panc89 | 0.018 | 0.190 | 0.018 | * |
| Panc1_Holo | Panc89_Holo | 0.073 | 0.290 | 0.073 | ns |
| Panc1_Holo | Panc89_Para | 0.016 | 0.190 | 0.016 | * |
| Panc1_Para | Panc89 | 0.018 | 0.190 | 0.018 | * |
| Panc1_Para | Panc89_Holo | 0.073 | 0.290 | 0.073 | ns |
| Panc1_Para | Panc89_Para | 0.016 | 0.190 | 0.016 | * |
| Panc89 | Panc89_Holo | 0.016 | 0.190 | 0.016 | * |
| Panc89 | Panc89_Para | 0.146 | 0.290 | 0.146 | ns |
| Panc89_Holo | Panc89_Para | 0.016 | 0.190 | 0.016 | * |
| Panc1 | Panc89 | 0.018 | 0.190 | 0.018 | * |
| Panc1 | Panc89_Holo | 0.073 | 0.290 | 0.073 | ns |
| Panc1 | Panc89_Para | 0.016 | 0.190 | 0.016 | * |

* *p*-value < 0.05, ** *p*-value < 0.01, *** *p*-value < 0.001, ns: *p*-value > 0.05

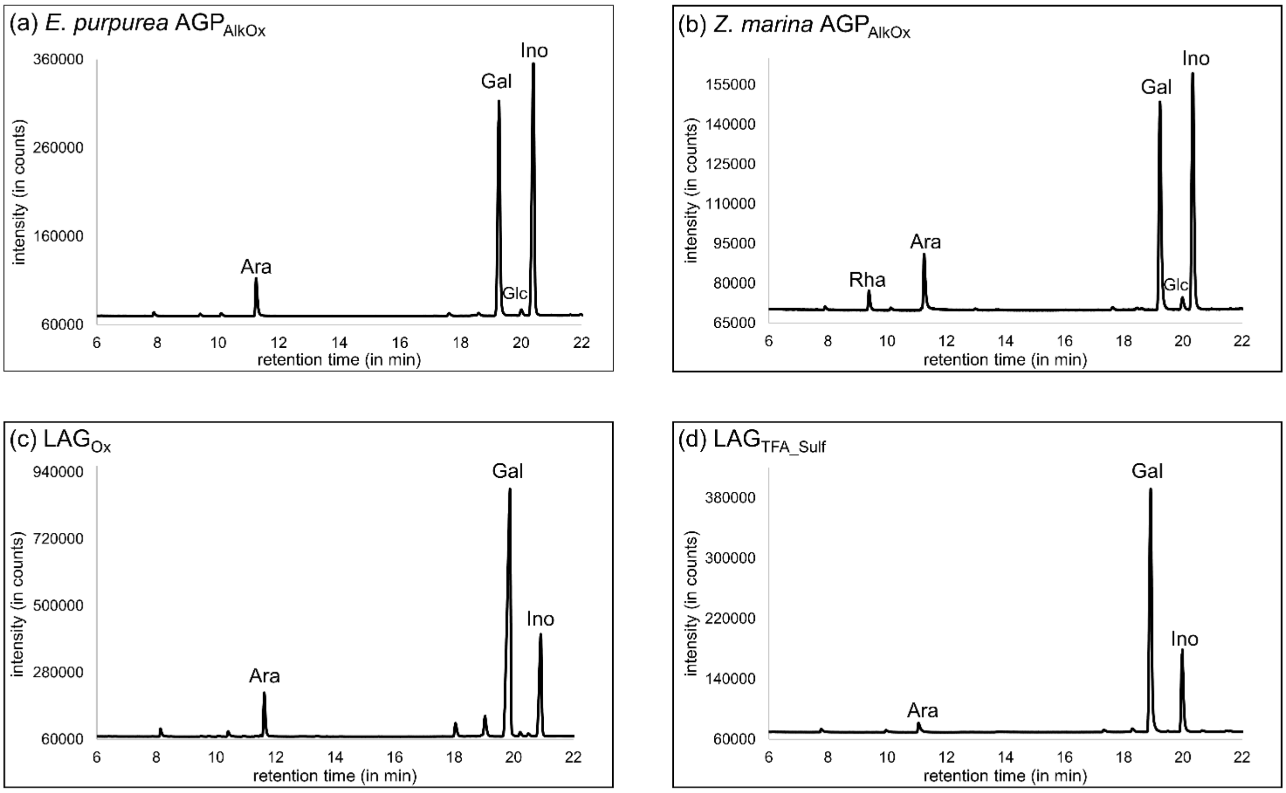


**Figure S1:** GC-FID chromatograms of the monosaccharide compositional analysis.

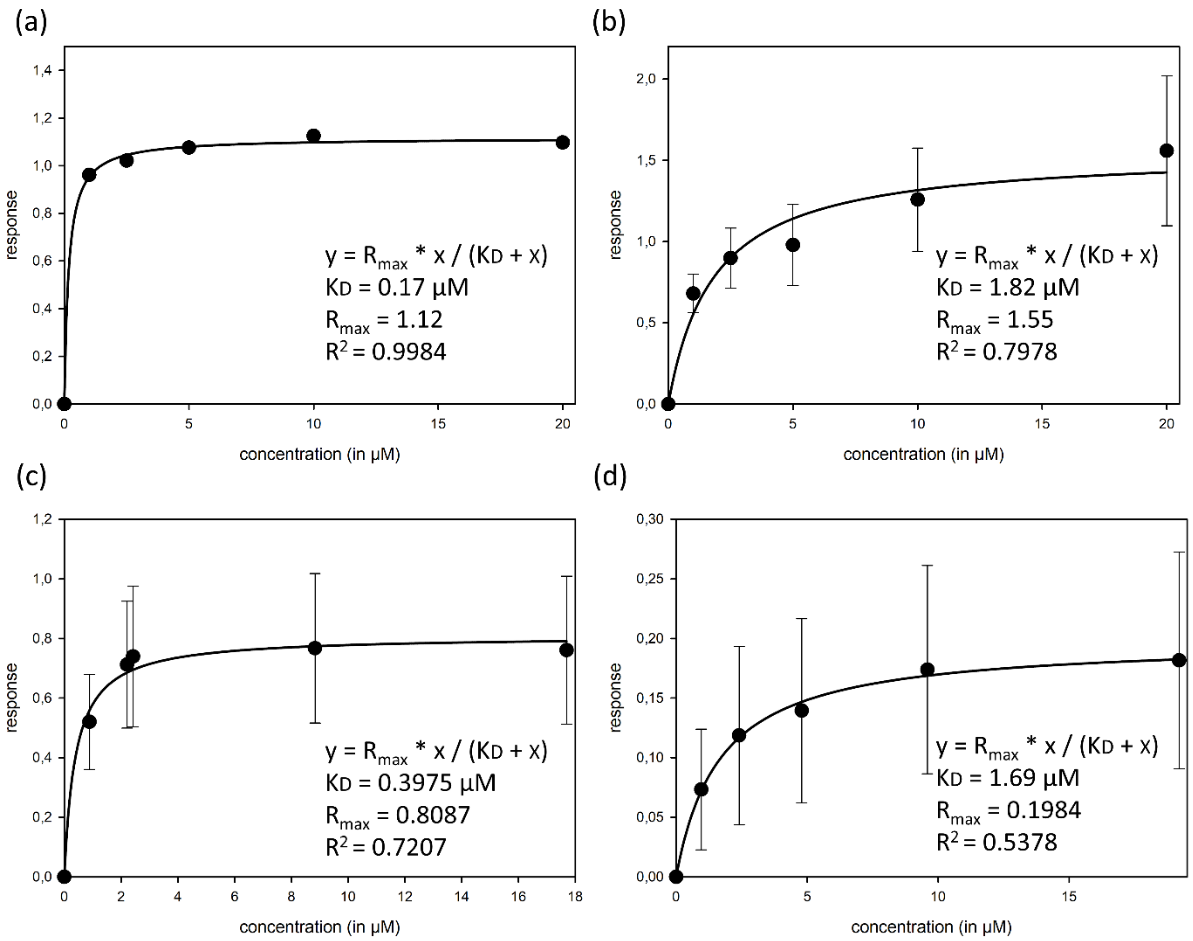


**Figure S2:** Results of the steady-state analysis for binding of (a) *Echinacea* galactan, (b) *Zostera* galactan, (c) $LAG_{TFA}$, and (d) $LAG_{TFA\_Sulf}$ to galectin-3

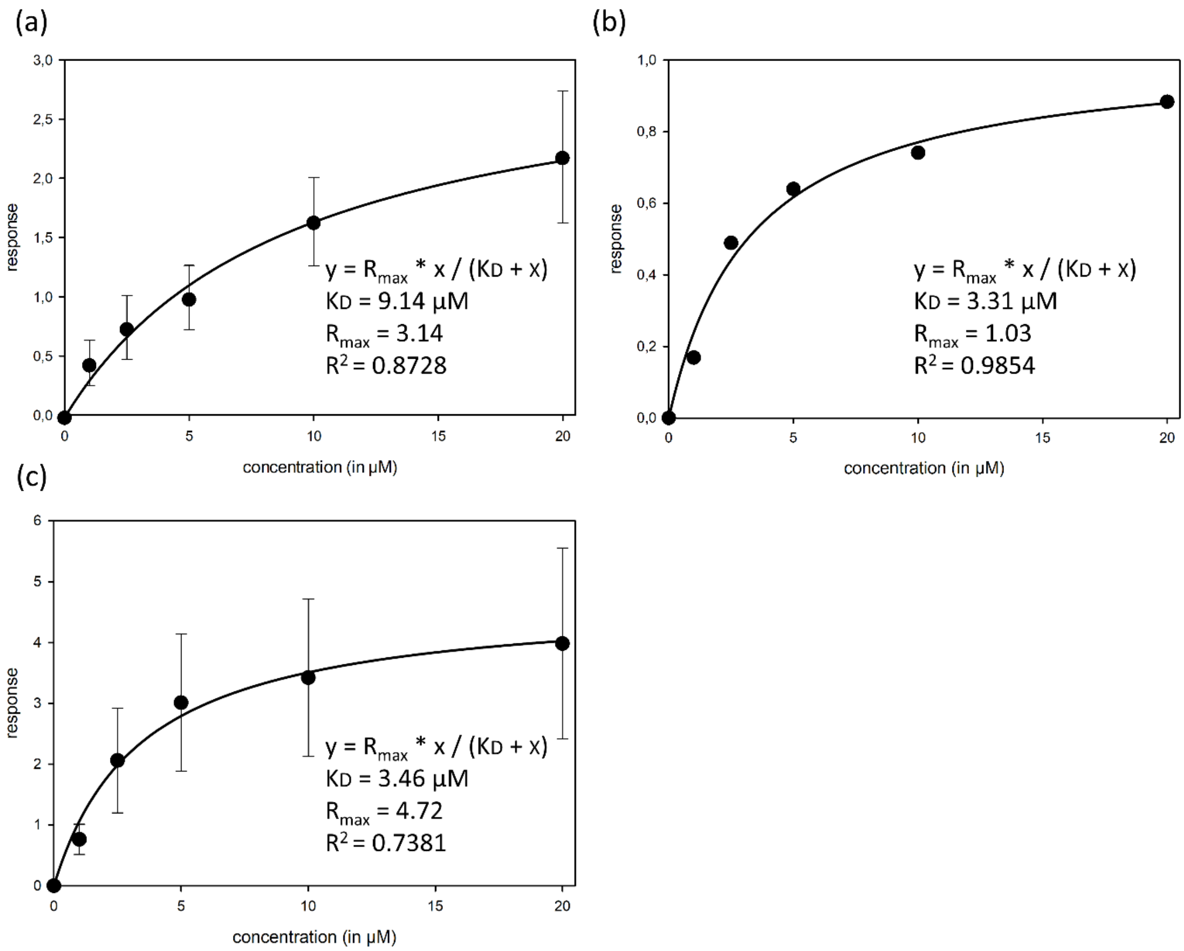


**Figure S3:** Results of the steady-state analysis for binding of different Yariv-reagents. (a) β-D-galactosyl-Yariv, (b) α-D-galactosyl-Yariv, and (c) β-D-lactose-Yariv to galectin-3.

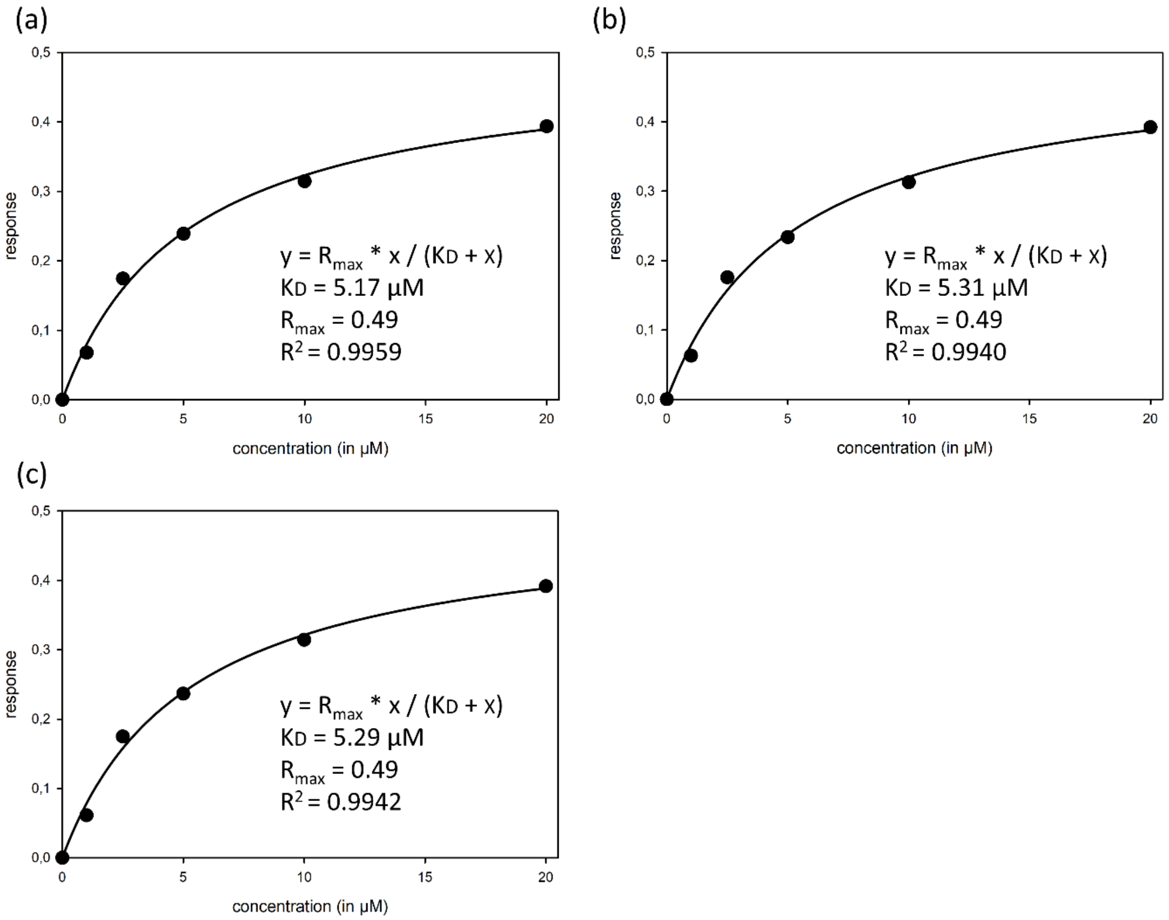


**Figure S4:** Results of the steady-state analysis for binding of three replicates of *Zostera* galactan (a-c) to galectin-9.

### References

[1] H. Knaack, L. Lenk, L.-M. Philipp, L. Miarka, S. Rahn, F. Viol, C. Hauser, J.-H. Egberts, J.-P. Gundlach, O. Will, S. Tiwari, W. Mikulits, U. Schumacher, J.G. Hengstler, S. Sebens, Liver metastasis of pancreatic cancer: the hepatic microenvironment impacts differentiation and self-renewal capacity of pancreatic ductal epithelial cells, Oncotarget 9 (2018) 31771–31786. https://doi.org/10.18632/oncotarget.25884.

[2] L.-M. Philipp, U.-U. Yesilyurt, A. Surrow, A. Künstner, A.-S. Mehdorn, C. Hauser, J.-P. Gundlach, O. Will, P. Hoffmann, L. Stahmer, S. Franzenburg, H. Knaack, U. Schumacher, H. Busch, S. Sebens, Epithelial and Mesenchymal-like Pancreatic Cancer Cells Exhibit Different Stem Cell Phenotypes Associated with Different Metastatic Propensities, Cancers 16 (2024). https://doi.org/10.3390/cancers16040686.
